# N-glycosylation blocks and simultaneously fosters different receptor-ligand binding sites: the chameleonic CD44–hyaluronan interaction

**DOI:** 10.1101/2020.03.23.004002

**Authors:** Joni Vuorio, Jana Škerlová, Milan Fábry, Václav Veverka, Ilpo Vattulainen, Pavlína Řezáčová, Hector Martinez-Seara

**Affiliations:** Department of Physics, University of Helsinki, P.O. Box 64, FI-00014 Helsinki, Finland; Computational Physics Laboratory, Tampere University, PO Box 692, FI-33014 Tampere, Finland; Institute of Molecular Genetics of the Czech Academy of Sciences, Videnska 1083, Prague 4, 142 20, Czech Republic; Institute of Organic Chemistry and Biochemistry of the Czech Academy of Sciences, Flemingovo nam. 2, Prague 6, 166 10, Czech Republic; Department of Cell Biology, Faculty of Science, Charles University, Vinicna 7, 128 00 Prague, Czech Republic; MEMPHYS – Centre for Biomembrane Physics

**Keywords:** CD44, hyaluronan, glycosylation, ligand-protein interaction

## Abstract

While DNA encodes protein structure, glycans provide a complementary layer of information to protein function. As a prime example of the significance of glycans, the ability of the cell surface receptor CD44 to bind its ligand, hyaluronan, is modulated by N-glycosylation. However, the details of this modulation remain unclear. Based on atomistic simulations and NMR, we provide evidence that CD44 has multiple distinct binding sites for hyaluronan, and that N-glycosylation modulates their respective roles. We find that non-glycosylated CD44 favors the canonical sub-micromolar binding site, while glycosylated CD44 binds hyaluronan with an entirely different micromolar binding site. Our findings show (for the first time) how glycosylation can alter receptor affinity by shielding specific regions of the host protein, thereby promoting weaker binding modes. The mechanism revealed in this work emphasizes the importance of glycosylation in protein function and poses a challenge for protein structure determination where glycosylation is usually neglected.

## Introduction

Glycosylation is a fundamental process where proteins are linked to complex oligosaccharides, glycans (1). Most of the proteins at the extracellular side of eukaryotic cells contain covalently linked glycans (2). Their structural roles include the mediation of interactions with the surrounding environment (3), facilitation of correct folding (4–6), and involvement in the assembly of membrane proteins (7), also by direct interaction with lipids (8). Glycans are also known to modulate the binding of ligands with several proteins, e.g., by masking the binding site (9–11). Such regulation is relevant, especially in most immune processes, such as activation and homing, guided by regulated remodeling of the glycans (12). However, the details of these modulation mechanisms are often poorly understood due to the glycans’ structural flexibility and dynamic nature (13, 14).

The transmembrane protein called CD44 is a key example of glycoproteins, whose functions are modulated by N-glycosylation (9, 15–19). Its primary task is to serve as a receptor for a carbohydrate polymer, hyaluronic acid (hyaluronan (HA)) (20, 21). This ligand–protein interaction mediates a variety of physiological processes such as white blood cell homing, healing of injuries, embryonic development, and controlled cell death (22). Recently, the CD44–HA interaction has also been utilized in the design of functional biomaterials (23). CD44 binds HA exclusively via its lectin-like hyaluronate binding domain (HABD). In the canonical form, CD44 is a 722 residue-long type I transmembrane protein from which HABD comprises the first 150 amino acids (20-169) after the signal peptide (24,25). Notably, human CD44-HABD contains five possible N-glycosylation sites (N25, N57, N100, N110, and N120) (24) (see Fig. 1) that are known to be occupied by highly branched N-glycans, especially in various cancer cell lines (18, 19, 26). The N-glycans elicit a dual effect on HA binding: while some glycan content favors the recognition of HA, the presence of negatively-charged sialic acids generally interferes or even blocks it (16, 27, 28). However, the molecular mechanisms underlying such a dual effect remain unclear. In fact, most of the currently available structural data of HA–CD44 complexes are derived from non-glycosylated constructs (24, 25, 29, 30), leaving the structural details of fully N-glycosylated HABD elusive.

**Fig. 1.**
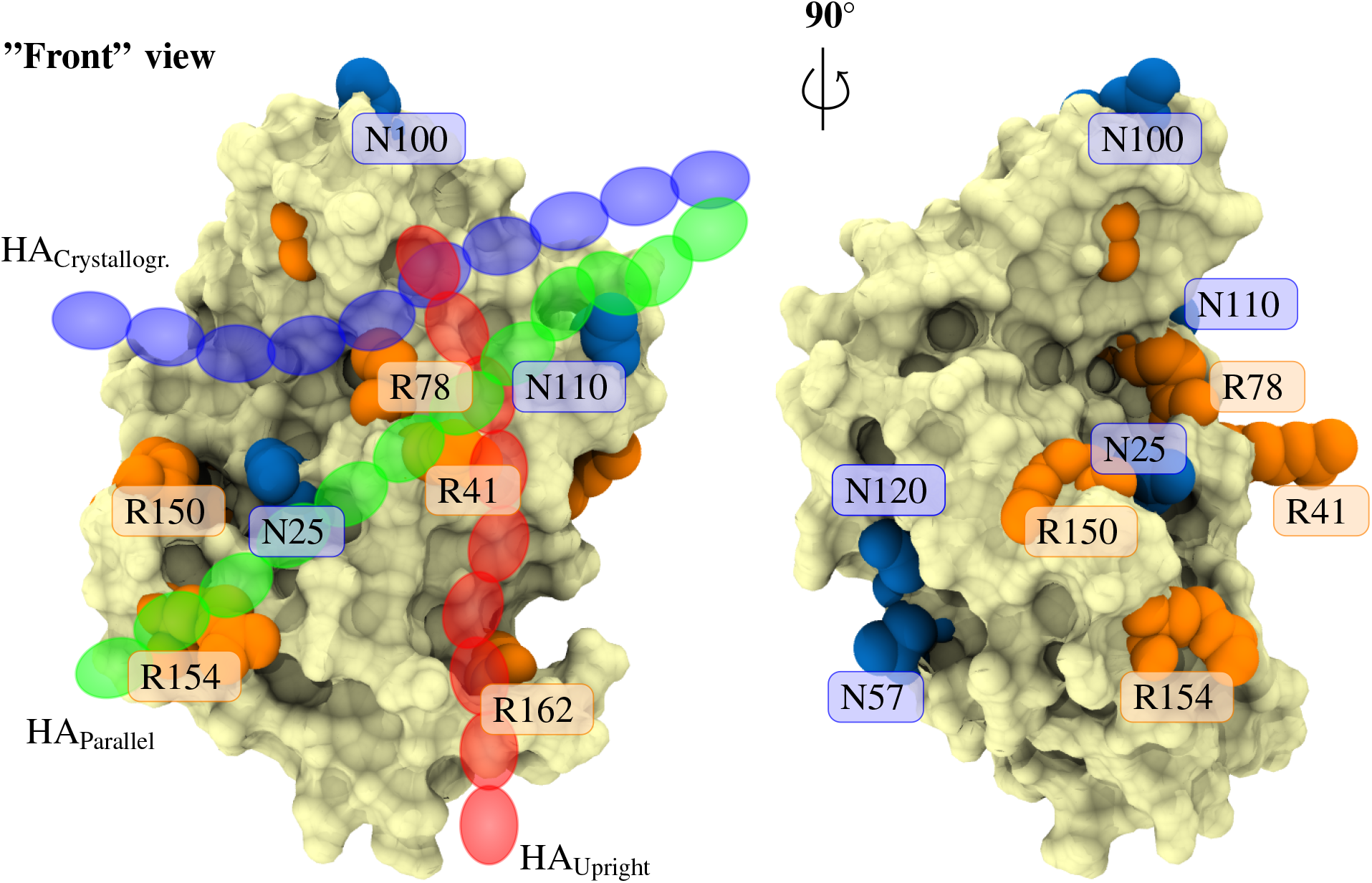
Human CD44-HABD (pale surface) with N-glycosylation sites (blue) and key HA binding arginines (orange) highlighted. Left panel also shows a schematic representation of HA in each binding mode (transparent blue, green, and red chains).

A shallow groove on the surface of the HABD forms the canonical binding site for HA. There the residue R41 stabilizes the binding in a pincer-like fashion (25, 31, 32). In addition to this so-called crystallographic binding mode (blue chain in Fig. 1), in our previous work, we postulated the existence of two potential lower-affinity binding modes called parallel (green chain in Fig. 1) and upright (red chain in Fig. 1) modes (14). These modes occupy the same general face of CD44-HABD, sharing to a large extent the R41-containing binding epitope. Additionally, each of these modes involves a second arginine residue that is distinct from that of the other binding poses (14). As a result, each mode covers a unique region of the CD44-HABD surface. Such separation of the binding sites allows their selective silencing via antibodies that target different regions of CD44 (33). It also suggests that the presence of N-glycosylations may affect each of the binding modes differently. This idea is the central hypothesis of this work.

In this study, we employed atomistic molecular dynamics (MD) simulations to unravel how complex N-glycans at N25, N100, and N110 cooperatively cover the canonical binding groove of CD44-HABD. This sugar shield hinders the accessibility and ligand availability of the canonical binding groove significantly, thereby promoting the secondary upright HA–CD44 binding mode over the crystallographic binding site. We then used NMR complemented by atomistic MD simulations to show that a few short HA oligomers can bind CD44-HABD simultaneously at distinct binding sites. The observed binding sites correspond to the previously characterized crystallographic (25), parallel, and upright binding modes (14). We further reveal that anti-CD44 antibody MEM-85 does not cross-block the canonical HA binding site in non-glycosylated CD44. Instead, it blocks HA binding to glycosylated CD44 (34, 35). These findings provide compelling evidence for the existence of a lower-affinity upright binding mode for HA. This binding mode overlaps with the binding site of MEM-85 and is promoted by N-glycosylation. The results demonstrate the existence of a new mechanism to control the ligand binding affinity of receptor proteins by promoting alternate binding sites by N-glycosylation.

## Materials and Methods

### A. Simulation system construction and models

We generated computational simulation models of glycosylated CD44-HABD. As the primary oligosaccharides, we employed fucosylated complex-type triantennary N-glycans, containing zero (asialo) or one (monosialo) terminal sialic acids per antenna, i.e., non-reducing termini. These oligosaccharide structures represent the predominant types in the so-called inducible (monosialo) hyaluronate binding phenotypes, together with a non-sialylated reference (asialo) (9,18, 26). To mimic the predominant CD44 glycovariants found recently in mouse myeloma cells (26), we glycosylated N25, N57, N100, and N110 with the above-described complex type N-glycans and N120 with a triantennary high-mannose type structure without fucosylation (Fig. 2c). We call these glycoforms *myeloma asialo* and *myeloma monosialo*, depending on the degree of sialylation. Additionally, to emulate the mutant proteins lacking the N25 and N120 glycans that also lead to the inducible phenotype (19), we constructed a monosialo glycoform, which lacks N-glycans at N25 and N120 (*partial monosialo*). Finally, we designed a fourth glycoform, where each of the five N-glycans is a charge-neutral core pentasaccharide (*full pentasaccharide*), to represent mildly glycosylated, less-cancerous cell types.

**Fig. 2.**
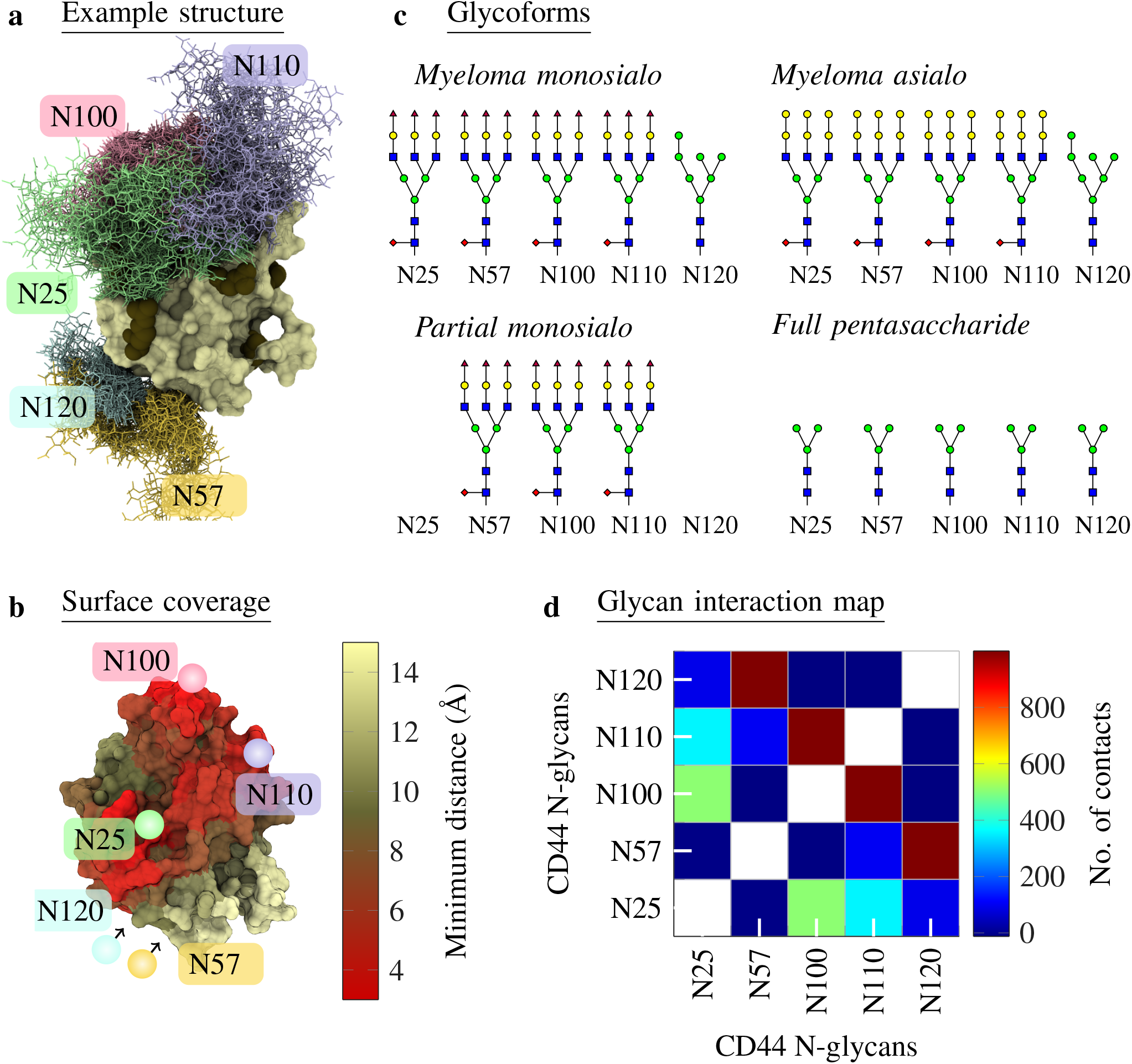
**a**: Example structure of CD44-HABD with the *myeloma monosialo* glycoform (system G2 in Table 1). CD44-HABD is colored pale, and different colors separate the glycans. Glycans are depicted at every 50 ns in a trajectory of 1000 ns. **b**: CD44-HABD with the surface colored according to the minimum distance to the N-glycans (not shown). Pale color corresponds to CD44–N-glycan distances over 15 Å, whereas bright red corresponds to distances less than 3 Å. The distance data have been averaged over 15 replicas (*myeloma monosialo* glycoform). **c**: Glycoforms used in this study. The symbols follow the Symbol Nomenclature for Graphical Representations of Glycans (50). **d**: Number of contacts (defined as distance < 0.6 nm) between the five N-glycans on CD44-HABD (*myeloma monosialo* glycoform). The results have been averaged over time and 15 replicas. Another simulation force field (CHARMM36) in Notes SC and SD shows consistent data.

We constructed the simulation systems using the crystal structure of human CD44-HABD (PDB:1UUH (36)). We then followed the steps described in our previous work to curate the 1UUH structure (14). This was followed by the *in silico* N-glycosylation of the HABD structure with the *doGly-cans* (37) tool. Before simulations, we inspected the readymade glycan structures visually (38) to confirm their correct configuration and stereochemistry. In every system, sodium and chloride ions were added to reach a typical physiological salt concentration of 150 mM, and to neutralize the charge of the system (Dang ions (39) for AMBER99SB-ILDN and default ions for CHARMM36 systems). The systems were solvated with the recommended TIP3P water model (40).

For each GLYCAM06-modeled system without a HA ligand (systems G1–4 in Table 1), we generated three different N-glycan starting configurations. Each configuration was used to start five replica simulations of 1000 ns, totalling to 15 replicas per glycoform. The CHARMM36 systems were simulated with three replicas (systems C1–2 in Table 1). Those three additional CHARMM36 systems had an added hyaluronate oligomer (18 monosaccharide units). We set them up initially to study HA binding to HABD, yet the oligomer never bound during the trajectories. Hence, we do not expect the hyaluronate to interfere with the folding of the N-glycans in those systems.

**Table 1.** List of simulated systems. *Glycoform* tells the N-glycan content on the CD44 HABD in each system. *HA* tells whether HA was present and which kind. *FF* lists the simulation force field. *Length* lists the duration of the simulations. Glycoforms of CD44 were experimentally found in: ^*a*^Ref. (26), ^*b*^Ref. (19), ^*c*^Ref. (9), ^*d*^Ref. (18), ^*e*^Ref. (24), ^*f*^ Ref. (16), and ^*g*^Ref. (17).

| System | Glycoform | HA | FF | Length (ns) | DOI DATA |
| --- | --- | --- | --- | --- | --- |
| G1<sup>a</sup> | Myeloma asialo | None | GLYCAM06 | 3 × 5 × 1000 | 10.5281/zenodo.3742147 |
| G2<sup>a</sup> | Myeloma monosialo | None | GLYCAM06 | 3 × 5 × 1000 | 10.5281/zenodo.3742149 |
| G3<sup>a,b</sup> | Partial monosialo | None | GLYCAM06 | 3 × 5 × 1000 | 10.5281/zenodo.3742154 |
| G4 | Full pentasaccharide | None | GLYCAM06 | 3 × 5 × 1000 | 10.5281/zenodo.3742158 |
| B1<sup>e</sup> | Non-glycosylated | HA<sub>18</sub> | GLYCAM06 | 8 × 1000 | 10.5281/zenodo.4005682 |
| B2<sup>f</sup> | Full GlcNAc | HA<sub>18</sub> | GLYCAM06 | 8 × 1000 | 10.5281/zenodo.4005689 |
| B3<sup>c,d,g</sup> | Full asialo | HA<sub>18</sub> | GLYCAM06 | 8 × 1000 | 10.5281/zenodo.4005691 |
| B4<sup>d,f</sup> | Full monosialo | HA<sub>18</sub> | GLYCAM06 | 8 × 1000 | 10.5281/zenodo.4005695 |
| B5<sup>b</sup> | Partial monosialo | HA<sub>18</sub> | GLYCAM06 | 8 × 1000 | 10.5281/zenodo.4005701 |
| B6<sup>f</sup> | Full polysialo | HA<sub>18</sub> | GLYCAM06 | 8 × 1000 | 10.5281/zenodo.4005707 |
| B7 | Full extended asialo | HA<sub>18</sub> | GLYCAM06 | 8 × 1000 | 10.5281/zenodo.4005740 |
| B8 | Full pentasaccharide | HA<sub>18</sub> | GLYCAM06 | 8 × 1000 | 10.5281/zenodo.4005743 |
| C1<sup>a,c,d</sup> | Myeloma asialo | HA<sub>18</sub> | CHARMM36 | 3 × 1000 | 10.5281/zenodo.3742175 |
| C2<sup>a,c,d</sup> | Myeloma monosialo | HA<sub>18</sub> | CHARMM36 | 3 × 1000 | 10.5281/zenodo.3742177 |
| G5 | Non-glycosylated | HA<sub>6</sub> | GLYCAM06 | 20 × 1000 | 10.5281/zenodo.3742160 |
| G6 | Non-glycosylated | HA<sub>6</sub> | GLYCAM06 | 10 × 1000 | 10.5281/zenodo.3742167 |

To understand how CD44 glycoprotein bind HA, we constructed GLYCAM06-modeled systems where HA_18_ was let to spontaneously form a complex with different glycoforms of HABD (simulations B1–8 in Table 1). In the initial frame, we positioned the HA oligomer to the water phase, roughly 2.5 nm away from the R41 residue (the most important binding residue). The reasoning behind this initial distancing is to avoid any bias in the binding, see Note SB. In total, we studied seven different glycoforms: *full asialo, full monosialo, partial monosialo, full extended asialo, full pentasaccharide, full GlcNAc*, and *full polysialo.* These glycoform names are further explained in the Note SA. Lastly, we used *nonglycosylated* HABD from Ref. 14 as reference system without glycosylation. For each glycoform, we performed eight replicas of 1000 ns, as listed in Table 1.

We also constructed an additional GLYCAM06-modeled system (20 replicas of 1000 ns) having non-glycosylated CD44-HABD together with three unbound (i.e., 1.5–2 nm from the protein surface as explained in Note SB) hyaluronate hexamers to study their spontaneous and simultaneous binding (system G5 in Table 1). That is, the carbohydrate fragments associated and/or dissociated from the protein spontaneously during the course of the simulation trajectories. Similarly, we generated systems (10 replicas of 1000 ns each) with CD44-HABD and two hyaluronate hexamers from which one was initially complexed to the crystallographic binding site, while the other was unbound (system G6 in Table 1). Tables S2 and S3 in Note SF list the observed association/dissociation cycles between the hexamers and HABD in simulations G5 and G6, showing that the sampling is adequate. All simulation data are publicly available in `zenodo.org`.

### B. Parameters for molecular dynamics simulations

Simulations were conducted using the GROMACS simulation software package (41). For every simulation, we employed the following protocol. First, to relax clashes produced in the building process, we performed a short energy minimization run with the steepest descent algorithm (1000 steps). Subsequently, we performed 1 and 2 ns equilibration runs in the NVT and NpT ensembles, respectively, with coordinates of the protein and glycans restrained. Finally, we conducted production runs of different lengths (see Table 1).

The production runs, along with equilibration, employed the leap-frog integrator with a time step of 2 fs. During the runs, periodic boundary conditions were used in all three directions, and the LINCS algorithm was used to keep all bonds constrained (42). Electrostatic interactions were treated with particle-mesh Ewald (PME) (43) with a cut-off of 1.0 nm for the real part. Lennard-Jones interactions were cut off at 1 nm. Neighbour searching for long-range interactions was carried out every ten steps. The V-rescale (44) thermostat was used to couple the systems to a heat bath of 310 K, while the Parrinello-Rahman (45) barostat was employed to keep the pressure at 1 bar. At the beginning of each production simulation, we assigned random initial velocities using the Boltzmann distribution at the target temperature. The CHARMM36 simulations used the default parameters provided by CHARMMGUI v1.7 (46). The simulation trajectories were saved every 100 ps. For other non-specified parameters, we refer to the GROMACS 4.6.7 (47, 48) defaults for the AMBER99SB-ILDN/GLYCAM06 systems or to the GROMACS 5.1.4 (41) defaults for the CHARMM36 systems.

### C. Analysis of simulations

All distances and numbers of contact were calculated with `gmx mindist` tool from the GROMACS 5.1.4 package, using a cutoff of 0.3 nm unless stated otherwise.

N-glycan coverage for a given binding mode, *C*^mode^, is calculated by comparing the interactions of HA with a nonglycosylated HABD to the interactions of N-glycans with their core HABD. The calculation averaged over each replica is conducted as follows:

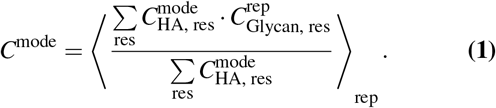

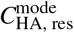 is the average coverage of the protein residue “res” by HA in a given binding “mode”. The parameter 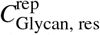 is the average coverage of the protein residue “res” by the N-glycans in each replica (“rep”). The average coverage (*C*_res_) is calculated as the ratio of frames where the distance between any atom of the HABD “res” to any atom of the target HA or N-glycans is closer than 0.3 nm.

### D. NMR spectroscopy

The proteins were expressed and purified as described in Ref. (33). The ^15^N/^1^H “heteronuclear single quantum coherence” (HSQC) spectra were acquired as described in Ref. (33) using a 350 μl sample containing 100 μM ^15^N-labeled CD44-HABD, a 350 μl sample containing 200 μM ^15^N-labeled CD44-HABD and 210, 415, and 620 μM hyaluronate hexamer (Contipro Group, Dolni Dobrouc, Czech Republic), using a 350 μl sample containing 100 μM ^15^N-labeled CD44-HABD and 200 μM unlabeled scFv MEM-85, or using a 320 μl sample containing 90 μM ^15^N-labeled CD44-HABD, 180 μM unlabeled scFv MEM-85, and 300 μM hyaluronate hexamer. The sequence-specific resonance assignment for free CD44-HABD was obtained as published in Ref. (33); signals could not be assigned for the following residues: Tyr42, Ser95, Asn100, Thr108, Ser109, Asn110, Ser112, Cys129 and prolines. The perturbations of ^15^N-labeled CD44-HABD signals in the HSQC spectra were monitored employing the minimal backbone chemical shift method (^15^N and ^1^H) (49).

## Results

### E. Complex N-glycans on CD44-HABD can cooperatively block its canonical binding site for hyaluronate

To characterize how N-glycans behave and fold on CD44-HABD, we *in silico* glycosylated a HABD structure (PDB:1UUH) with *myeloma asialo, myeloma monosialo, partial monosialo*, and *full pentasaccharide* N-glycan profiles (Systems G1–4 in Table 1 depicted in Fig. 2c). We then simulated each glycoform through 15 replicas. An average minimum distance between the complex N-glycans and the protein, as mapped onto the surface of HABD (Fig. 2b), reveals that in the *myeloma* glycoforms, the N-glycans cover a significant fraction of the protein surface. That is, with the complex oligosaccharides in *myeloma monosialo* and *myeloma asialo* glycoforms, the N25 glycan can interact intimately with the nearby N100 and N110 glycans, forming a sugar shield that covers the canonical binding site of hyaluronate (Fig. 2a). Furthermore, the contact map for the five N-glycans in the *myeloma monosialo* glycoform (Fig. 2d) shows the glycans at N25, N100, and N110 to establish, on average, several hundred intermolecular contacts, which are possible only if the three N-glycans become interconnected in the region that resides over the crystallographic hyaluronate binding groove. These results clearly show how complex N-glycans, facilitated by inter-N-glycan interactions, shield a significant portion of the hyaluronate binding face of HABD.

To study the spontaneous binding of HA to the N-glycosylated CD44-HABD, we performed simulations where both molecules were initially significantly separated. In this setting, the molecules can interact in a spontaneous manner without any apparent bias. These simulations refer to sets B1–8 in Table 1. Typical binding complexes arising from this set-up are shown in Fig. S8a–d in Note SH.

Comparing the final (*t* = 1000 ns) HA–HABD and HA– N-glycans interface areas after the spontaneous binding of HA to the glycosylated HABD (Fig. S8e in Note SH) reveals how the glycans hinder the recognition and how different glycoforms influence this process. While the HA–N-glycans interface obtains average values of 6–10 nm^2^ with all glycoforms larger than *full GlcNAc*, the HA–CD44 interface varies significantly depending on the N-glycan content. The shortest *full GlcNAc* glycoform displays HA–CD44 binding similar to that of the non-glycosylated reference with recognizable binding modes, showing that these neutral sugar units do not obstruct the ligand binding. Instead, they offer more binding surface for HA compared to non-gkycosylated HABD. Medium-sized neutral glycans (i.e., *full pentasaccharide* and *full asialo)* display HA–HABD interfaces (~6 nm^2^) slightly lower than the non-gkycosylated reference (~8 nm^2^). While these glycoforms also provide additional interaction sites for HA through the larger size of the N-glycans, they also cover the important binding residues, preventing the formation of clear HA–CD44 binding modes. Agreeing with previous experimental findings, the sialylated glycoforms (*full monosialo*, *partial monosialo*, and *full polysialo*) display relatively low HA-HABD interfaces, with the partially glycosylated form showing the strongest binding to the protein. Together these results indicate that both the size of the N-glycans and charge (number of sialic acids) abrogate the binding of HA.

### F. N-glycans foster the occupancy of a secondary hyaluronate–CD44 binding mode

Table 2 lists the coverage of each of the three binding sites by the N-glycans. In the tested glycoforms (systems G1-4 in Table 1), amino acid residues distinct to the CD44-HABD binding modes exhibit a coverage of about 20 to 50 %. The somewhat high standard errors indicate a large replica-to-replica variance in the folding of the N-glycans, as well as slow interconversion between the folding patterns. The use of 15 replicas, however, ensures a reasonable sampling of the possible patterns. In all cases, the crystallographic binding site is most significantly obstructed by the N-glycans, while the upright site is obstructed the least. Furthermore, coverage values calculated for the key hyaluronate binding residues of CD44-HABD (see Note SE) reveal how the key residues that are specific to the upright mode, such as K38 and R162, are generally less covered by the N-glycans. These observations imply that the lower-affinity upright mode is the most accessible binding configuration in a glycosylated CD44-HABD. Strikingly, we observe minimal differences between the *myeloma monosialo* and *myeloma asialo* glycoforms, where the oligosaccharides are of the same length. However, the coverage values decrease notably with reduced glycan content in the *partial monosialo* or the shorter *full pentasaccharide* glycoforms. Like the myeloma-derived CD44-HABDs, the *partial monosialo* glycoform also displays a large number of contacts between the glycans N100 and N110 (Note SD). Their interaction is, however, less prone to disturb the crystallographic binding site as the N25-linked glycan is missing (Note SD). The *full pentasaccharide* glycoform is fully glycosylated but entails shorter oligosaccharides, which therefore limit the degree of protein coverage. These observations suggest that it is predominantly the degree of glycosylation and the size of the attached oligosaccharides that determine the coverage of the binding site. The inclusion of sialic acids has little effect on the coverage when compared to similarsized non-sialylated N-glycans.

**Table 2.** Total N-glycan coverage of the residues involved in each binding mode. Data are calculated from systems G1–4 in Table 1. The contributions of each residue to each binding mode are extracted from our previous work (14). The results indicate how much of the CD44-HABD surface (that is critical to hyaluronate binding) is covered by N-glycans.

| Binding mode | Realistic asialo: coverage (%) | Realistic monosialo: coverage (%) | Partial monosialo: coverage (%) | Pentasaccharides: coverage (%) |
| --- | --- | --- | --- | --- |
| Cryst. | 50 ± 13 | 51 ± 8 | 35 ± 18 | 32 ± 7 |
| Parallel | 48 ± 14 | 46 ± 11 | 34 ± 17 | 26 ± 6 |
| Upright | 29 ± 12 | 30 ± 7 | 21 ± 13 | 16 ± 4 |

Figure S9 in Note SH compiled from the spontaneous binding simulations shows that the *non-gkycosylated* HABD expresses the most interactions between HA and the arginines at the crystallographic binding groove (R41 and R78) compared to all the glycosylated HABDs. This indicates that the presence of N-glycans generally decreases the accessibility of these key HA binding residues. Consistently, the flanking arginines (R150, R154, and R162) are relatively more prone to interact with the ligand in the glycosylated cases, further suggesting that the binding modes involving these flanking arginines are activated in the glycosylated receptor. For additional observations from the spontaneous binding simulations, see Note SI.

### G. Antibody MEM-85 does not cross-block hyaluronate binding to non-glycosylated CD44

scFv MEM-85 antibody prevents the binding of hyaluronate to glycosylated CD44 (34, 35). We used NMR to probe whether the same antibody prevents hyaluronate from binding a non-glycosylated CD44-HABD. The hyaluronate and antibody induced changes are clearly visualized in an overlay of the ^15^N/^1^H HSQC spectra for the free ^15^N-CD44-HABD, ^15^N-CD44-HABD in complex with either hyaluronate hexamer (in a 3-fold molar excess), or scFv MEM-85 (in a 2-fold molar excess), and both scFv MEM-85 (in a 2-fold molar excess) and hyaluronate hexamer (in a 3-fold molar excess), see Fig. 3a. The observed changes can be interpreted as local perturbations/contacts in the vicinity of a given residue but they may also reflect a non-local secondary perturbation of some sort.

**Fig. 3.**
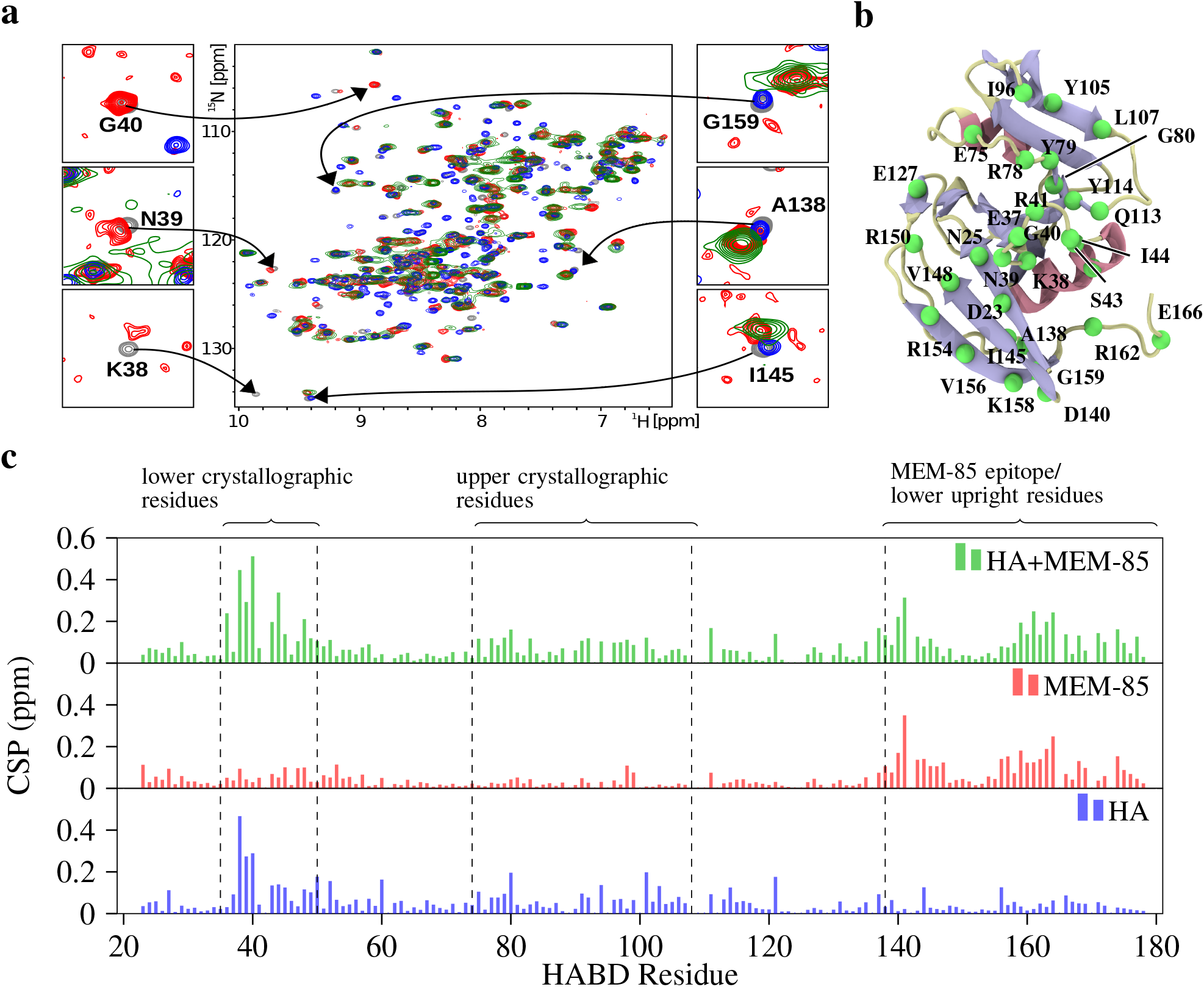
NMR confirms the simultaneous binding of scFv MEM-85 and hyaluronate to CD44-HABD. **a**: 2D ^15^N/^1^ H HSQC spectra are shown for free ^15^N-CD44-HABD (grey), ^15^N-CD44-HABD in complexes with hyaluronate hexamer (3-fold molar excess; blue), scFv MEM-85 (2-fold molar excess; red), and both scFv MEM-85 (2-fold molar excess) and hyaluronate hexamer (3-fold molar excess; green). Details of the signals are shown for selected residues from the hyaluronate-perturbed (left) and antibody-perturbed (right) regions of CD44-HABD. **b**: Illustration of the CD44-HABD (PDB:1UUH), highlighting the residues mentioned in the main text. From the residues, only the C_α_ atom is depicted (green). Coloring of the protein is based on secondary structure, such that coils are pale, sheets are blue, and helices are red. **c**: Histograms of the minimal combined chemical shift perturbation (CSP) versus the protein sequence are shown for ^15^N-CD44-HABD in complex with hyaluronate hexamer (blue), scFv MEM-85 (red), and both scFv MEM-85 and hyaluronate hexamer (green).

Residues from the hyaluronate-perturbed region such as K38, N39, and G40 exhibit similar spectral behaviour for the mixture of hyaluronate and antibody as for hyaluronate alone, i.e., their signals disappear. On the other hand, residues from the antibody-perturbed region (33) such as A138, I145, and G159 —necessary for upright mode— exhibit similar perturbations for the complex with both hyaluronate and the antibody, as in that of the antibody alone. Moreover, we calculated the histograms of the minimal combined chemical shift perturbation with respect to the free CD44-HABD spectra along its sequence (Fig. 3c). The obtained chemical shifts indicate that the spectra of the complex of CD44-HABD with both the antibody and hyaluronate still possesses the antibody-induced changes (residues within mainly the C-terminal segment of CD44-HABD) in addition to the hyaluronate-induced changes (residues within mainly the N-terminal segment of CD44-HABD). This clearly suggests the simultaneous binding of both hyaluronate hexamer and scFv MEM-85 to the non-glycosylated recombinant CD44-HABD.

In addition, the signals in the spectrum obtained for ^15^N-CD44-HABD in the presence of both antibody and hyaluronate hexamer are significantly broadened relatively to the signals in the spectra obtained for binary mixtures, as expected for a higher molecular weight of the ternary complex.

### H. Short hyaluronate oligomers bind to CD44-HABD simultaneously at distinct binding sites

We analyzed the individual signals in the HSQC spectra for ^15^N-CD44-HABD titrated with hyaluronate hexamer; signals located in crowded areas of the spectra, including R41, were not taken into account to avoid ambiguity. This analysis revealed two trends (Fig. 4). Certain backbone amide group signals exhibited an instant shift or disappearance already at the hyaluronate to CD44-HABD ratio of 1:1, which indicates a strong interaction in the sub-μM range of the respective residues with hyaluronate, while other signals shifted gradually during the individual titration steps, suggesting a relatively weaker interaction (>10 μM) (Fig. 4a). In addition, several signals exhibited doubling, connected either with an instant shift or a gradual shift (Fig. 4b). This points out residues which interact only in a fraction of CD44-HABD molecules with hyaluronate, and/or interact in two different modes. Specifically, the signals of the following residues exhibited instant disappearance: K38, G40, G80, Y114; in-stants shift: S43, I44, Y79; instant shift with doubling: D140, R150, R154, V156, T174; gradual shift: D23, N25, E37, E75, I96, Y105, Q113, E127, V148, G159, R162, E166; and gradual shift with doubling: N39, R78, L107, K158, N164, D175.

**Fig. 4.**
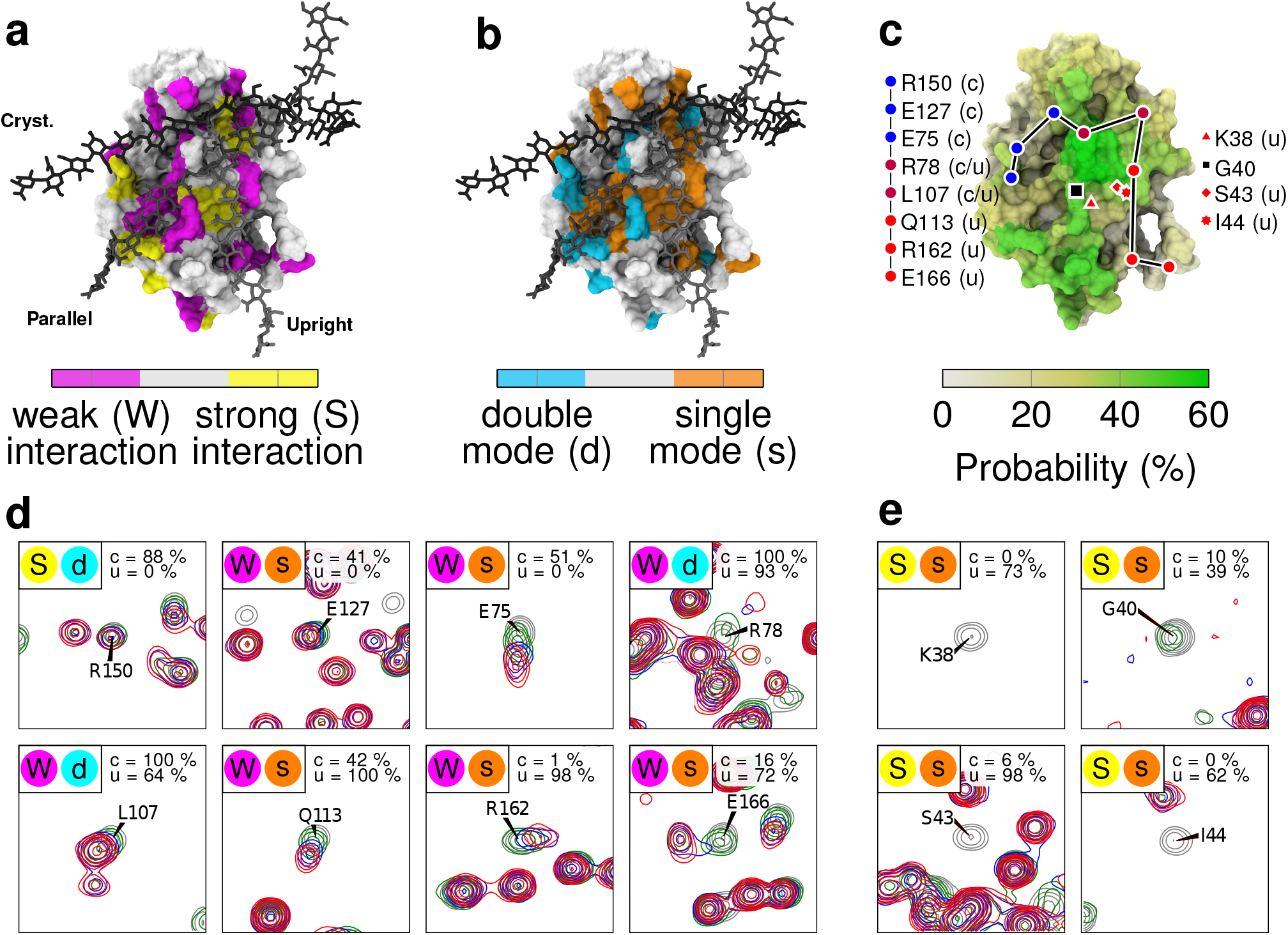
NMR titration of CD44-HABD by hyaluronate hexamer. Structures shown in a, b, and c are extracted from our previous study, where the CD44 coordinates are based on PDB ID 1UUH. **a**: Selected ^15^N/^1^ H HSQC signals are sorted in two groups — residues with an immediate shift/disappearance of the signal, i.e., strong interaction (yellow color), and residues with a gradual shift, i.e., weak interaction (magenta color). **b**: The signals are sorted in two groups — residues with a single signal, i.e., single binding mode (orange color), and residues with a doubled signal, i.e., double binding mode (cyan color). Hyaluronate hexadecamers are shown in three possible binding modes distinguished by shades of gray – the crystallographic (*cryst*.), parallel (*parallel*), and upright (*upright*) mode. **c**: Hyaluronate-perturbed residues in simulations. The colored surface displays the probability of a given residue to be in contact with HA6 in our simulations (G5 in Table 1). Filled circles highlight the positions of selected residues from both the crystallographic and upright modes, which were perturbed by hyaluronate binding in our NMR experiments (cf. panel d). These marks are also colored based on the predominant binding mode of the highlighted residues in our earlier simulations (14). Lines between the residues are drawn to guide the reader. The position of residues belonging to the R41-containing binding epitope is also shown (cf. panel e). Fig. S5 in Note SF shows similar surface data for both hexamer systems (Systems G5 and G6 in Table 1) **d e**: Selected ^15^N/^1^ H HSQC signals are shown for free ^15^N-CD44-HABD (grey), ^15^N-CD44-HABD with equimolar hyaluronate hexamer (green) and two-fold (blue) and a three-fold (red) molar excess of hyaluronate hexamer. Colored circles (top left) indicate to which category the residue falls in panels a and b. Probability of a given residue to interact with HA in each binding mode in our earlier simulations (14) is indicated in the top right corner of each graph.

Next, we mapped the critical residues involved in either strong or weak interaction with hyaluronate (Fig. 4c) and the residues interacting with hyaluronate in a single/double mode (Fig. 4d–e) onto the surface of a computational model of CD44-HABD (residues 20-169) (14). This illustrates that the surface patches associated with all the three modes are affected in our hyaluronate titration experiments. Notably, the linear patch including residues K38, S43, I44, Y79, G80, Y105, Q113, Y114, R162, and E166 outlines the binding site for the upright binding mode. Moreover, the doubling of the signals in the C-terminal portion of CD44-HABD (residues D140, R150, R154, V156, K158, and N164) indicates the coexistence of the parallel and upright modes with the crystallographic mode. The NMR data, therefore, demonstrate that the short hyaluronate hexamer can, especially in higher molar excess, bind to non-glycosylated recombinant CD44-HABD simultaneously in several modes at distinct binding sites.

To further explore the simultaneous binding of hyaluronate on CD44-HABD, we performed a set of MD simulations with three hyaluronate hexamers binding to CD44-HABD (simulation G5 in Table 1). In these systems, the hyaluronate hexamers are initially in an unbound state (see Note SB), and thus, readily able to sample the space and find their respective binding sites during the course of the simulations (See Table S2 in Note SF). Fig. 4c shows the probability of the HABD surface to be in contact with HA, which correlates with the combined chemical shift perturbations observed in NMR. Additionally, Fig. S6 in Note SF shows a contact profile similar to the chemical shift profile recorded in NMR (Fig. 3), indicating that our experimental and computational results are in agreement.

## Discussion

We employed atomistic MD simulations and NMR to shed light on ligand-receptor interactions of CD44 and hyaluronate to unravel how N-glycosylation modulates the interactions. MD simulations showed that in the crystallographic mode (sub-μM), N-glycans on CD44-HABD collectively shield the primary binding residues for hyaluronate. The shielding effect in this canonical binding mode is the strongest when complex type N-glycans occupy the N-glycosylation sites N25, N100, and N110. They are the most typical oligosaccharides found in these N-glycosylation sites (26) and are sufficiently long to interlock over the canonical hyaluronate binding groove, thereby severely hindering its availability for the ligand.

Backing these observations, our HA binding simulations with glycosylated HABD show how the smaller N-glycan types, such as simple GlcNAc residue, lack both the reach and charge necessary to influence the binding of HA in a negative way. Instead, the presence of GlcNAc residues offers additional binding surface for HA, thereby possibly advocating the recognition of HA by providing additional polar interaction sites and minimal hindrance to the binding. This observation is in line with previous research that has shown with metabolic glycosidase enzymes that GlcNAc residues on CD44-HABD have a positive effect on the binding of HA (16). Our simulations also show that once the size of the glycans on HABD increase, they start to prevent the entry of the ligand into its main binding site. High concentration of sialic acids further amplifies this effect through the increased size and negative charge of the glycans. Overall, our results from the spontaneous binding og HA to glycosylated HABD are in good qualitative agreement with previous experiments that have assessed the effect of different glycoforms, showing similar N-glycan-related size and charge-dependence for the binding of HA (9, 16, 27).

We also found that the N-glycosylation of CD44-HABD promotes a secondary, less shielded but weaker (>10 μM) hyaluronate binding site, which corresponds to the upright binding mode characterized previously by us (14) and also suggested by others (24). The results also revealed the degree of glycosylation and the size of the attached oligosaccharides to be the key factors in determining the coverage of the binding site, while the inclusion of single sialic acids to the glycan termini was found to have only a minor additional effect when glycans of equal length were compared to one another. Thus, it can be speculated, in the case of CD44– HA binding, that the binding-inhibiting role of monosialic acids stems from the more extended nature of the oligosaccharides and the resulting increase in the degree of coverage. The negative charge may play a more significant role in the case of polysialylated sugars, see Note SH. Furthermore, if the glycosylation site N25 lacks sufficiently long glycans, the propensity to interlock with the N100 and N110 glycans decreases, thereby substantially decreasing the coverage of the crystallographic site, resulting in a more exposed site to the ligand. This is again in line with findings that have suggested some glycosylation patterns do not decrease the hyaluronate binding (16).

Our NMR experiments support the notion of distinct hyaluronate binding sites on non-glycosylated CD44-HABD, which provides substantial evidence for the existence of separate hyaluronate binding modes. Strikingly, the residues perturbed in NMR match closely to those involved in the crystallographic, parallel, and upright binding modes. In our previous computational work, we illustrated the dynamic nature of the HABD–HA interactions outside the R41 epitope, especially in the case of the crystallographic binding mode (see Note SG). Similarly, the strong versus weak combined chemical shift perturbations in Fig. 4a show both the R41 epitope and upright groove to give predominantly strong interaction signals, while other regions flanking the R41 epitope tend to give out weak interaction signals, corresponding with the increased mobility of the bound HA in those regions. Despite the dynamic interactions, the importance of such weak binding sites to the overall strength of the binding is found to be high in a related protein–carbohydrate interaction (51). The dynamics of the bound HA can be visualized in Note SG.

The experimental results also agree well with the findings of our simulations of multiple hyaluronate hexamers with CD44-HABD, showing a similar hyaluronate–HABD binding profile (Fig. S6 in Note SF). The NMR readouts also show that the anti-CD44 antibody MEM-85 co-binds with hyaluronate on a non-glycosylated CD44, thus having a minimal effect on hyaluronate binding in this case. Conversely, the literature clearly states that MEM-85 blocks the hyaluronate binding of a glycosylated CD44 (34, 35), implying the existence of a lower-affinity binding mode, whose binding site overlaps with the binding site of MEM-85. The MEM-85 epitope is known to be located around the residues Glu160, Tyr161, and Thr163 (33). As these residues are also a part of the upright mode, our results hint towards the existence of such binding.

Providing further evidence for the existence of the upright mode, when CD44-Ig (immunoglobulin) fusion proteins were expressed in COS cells and hence were presumably glycosylated, both MEM-85 and hyaluronate binding were significantly reduced by the mutation of K38 to arginine (34). According to our previous work, K38 is exclusive to the upright mode (14), which further implies that glycosylated CD44 favors to bind hyaluronate with the upright mode over the canonical crystallographic binding. We also note that distinct N-glycosylation profiles, e.g., ones that include an increasing amount of sialic acids, might cause different alterations to the binding.

CD44–HA interaction is known to display glycosylation-dependent levels of activation (9) and binding affinities (16). The activation levels have been attributed to varying degrees of sialylation (19, 52), yet the glycosylation dependent binding affinities could stem from the simultaneous masking of high-affinity binding sites and promotion of secondary sites. Such activation-dependent regulation of glycan remodeling is undoubtedly known to be a major mechanism driving cell motility, e.g., in the immune response (12). CD44, in particular, is a hyaluronate-dependent leukocyte homing receptor that mediates both rolling interactions (53) and cellular transmigration (54). In such processes, tightly regulated affinity is required to enable dynamic velcro-like interactions between leukocytes and endothelial cells at inflamed tissue.

It is known that glycans stabilize or promote specific protein conformations (4–6), dimer interfaces (55), or orientations (8, 56), which ultimately affect ligand binding. There is also evidence of oligosaccharides that mask and shield specific parts of the protein surface (11, 57). N-glycosylations are also generally quite well known to protect large regions of the protein surface from, e.g., non-specific interactions or proteolytic cleavage (58). The novelty of the present work lies in the fact that, in addition to all these features, N-glycosylation has an extremely valuable and hitherto unknown mechanism of action: N-glycosylation can control the affinity of ligand–receptor interaction by selectively blocking binding sites and promoting others.

## Supporting information

Rebuttal elife

## ACKNOWLEDGEMENTS

HMS acknowledges support from the Czech Science Foundation (19-19561S). I. V. acknowledges financial support from the Academy of Finland Center of Excellence program, Sigrid Juselius Foundation, and the European Research Council (CROWDED-PRO-LIPIDS). JŠ, MF, VV, PŘ acknowledge funding from projects RVO 61388963 and 68378050 awarded by the Academy of Sciences of the Czech Republic and by the Ministry of Education of the Czech Republic, projects LO1304 (program ‘NPU I’) and CZ.02.1.01/0.0/0.0/16_019/0000729 (program OP RDE). We also acknowledge CSC-IT Center for Science (Espoo, Finland) for providing the computing resources that rendered this work possible.

## Supplementary Note 1: Additional methods and results

### SA. Studied CD44 glycoforms

The N-glycan types used in this study are presented in Figure S1. Using these oligosaccharide types, we constructed several glycoforms of CD44 HABD, as listed in Table 1. The naming of each glycoform is composed of two parts (excluding the nonoglycosylated reference named as *non-gkycosylated*): the first part signifies the degree of glycosylation and second part refers to the nature of the oligosaccharides used. In the first part of the name, ‘full’ refers to all the five N-glycosylation sites being occupied by N-glycan, while ‘partial’ means only sites N57, N100, and N110 are N-glycosylated. Term ‘myeloma’ refers to a glycoform where site N120 is glycosylated with high mannose type of oligosaccharides, while the remaining sites are N-glycosylated with the type of oligosaccharides given in the latter part of the name.

**Fig. S1.**
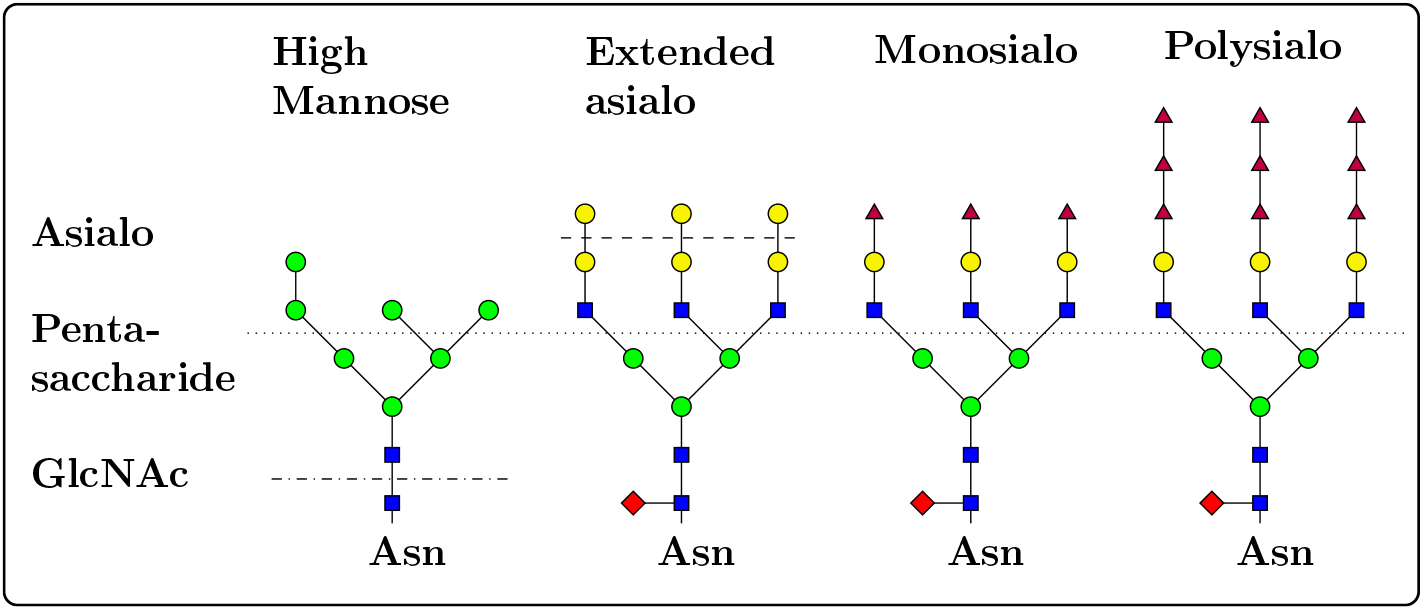
N-glycan structures employed for *in silico* glycosylation of the CD44 HABD. The horizontal dash dotted line separates the pentasaccharide core (below) from the antennae (above), while the short dashed line separates extended asialo N-glycan (above) from a shorter one (below). The dotted line separates the core pentasaccharide (below) from the antennae (above). Glycans are drawn according to the CFG symbol nomenclature.

The studied glycoforms represent a range of differently-sized CD44 glycoproteins, modeling the N-glycan structures presumably present in the inactive (*full polysialo*), inducible (*full monosialo, partial monosialo* (19)), and active (*full asialo, full GlcNAc*) HA binding phenotypes, having varying sialic acid content and size (9, 18). *Full pentasaccharide* and *full extended asialo* glycoforms were generated to further evaluate the the effect of N-glycan size. The *myeloma* glycoforms mimic the predominant galactose-terminating and sialic acid-terminating CD44 glycovariants found recently in the mouse myeloma cells (26). Non-glycosylated HABDs serve as a reference to the glycosylated proteins.

### SB. Avoiding simulation bias in the observed binding modes

When setting up any simulation system, one can bias the obtained results. To avoid favoring any particular binding mode between HA and HABD-CD44, the following two necessary steps were followed. First, we performed several replicas for each studied glycoform (Table 1). Second, HA was placed away from any potential binding site (1.5-2.5 nm). Here, we provide an example of such placing for the hyaluronate hexamers in simulations, see Fig. S2. This sort of placement reduces the risk of biasing HA’s binding to any of the studied binding sites substantially. Simultaneously, the initial proximity enhances the chances of finding the binding modes in the allocated simulation time.

### SC. Force field comparison

All the critical simulations were repeated with the CHARMM36 force field (systems C1–2 in Table 1), and their conclusions were consistent with the combination of AMBER99SB-ILDN and GLYCAM06 discussing along the text. In this manner we can assure that the results are not force field-dependent.

Table S1 lists the coverage of each HA binding mode by the glycans using the CHARMM36 force field (simulations C1-2) in Table 1. These values are in general around 15 to 25 % smaller than that of the GLYCAM06 systems (see Table 1 in the main text). Yet, they also share the same trend: the crystallographic and parallel modes are the most and upright mode the least covered. In the CHARMM36 systems, the replica-to-replica variances are in general smaller than for GLYCAM06 as the N-glycans show greater mobility.

### SD. N-glycan versus N-glycan contacts

Figure S3 shows numbers of contacts between the five N-glycans on all tested glycoforms and force fields.

### SE. Interaction of hyaluronate binding residues with N-glycans

Figure S4 presents a contact histogram of the most prominent binding residues found from the literature (14). It shows the binding of these residues to the N-glycans of each N-glycosylation site. For reference, the histogram also shows how these residues bind to hyaluronate oligomers (HA_16_ in our previous study (14).

**Fig. S2.**
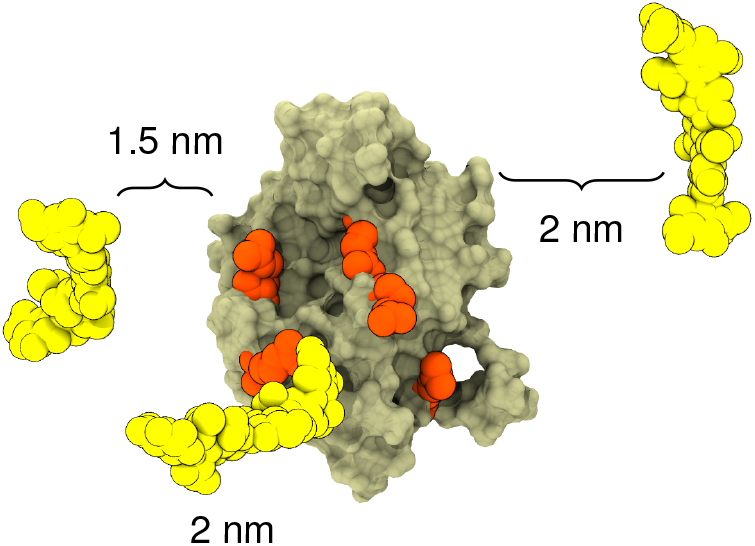
Representative starting configuration in System G5 simulations. Pale surface represents HABD; orange spheres are arginines R41, R78, R150, R154, and R162 (See Fig. 1; and yellow spheres are the HA hexamers. Each hexamer was placed randomly 1.5-2.0 nm away from the protein surface. This initial configuration quaranteed that the hexamers were able to sample the protein surface freely during the course of the simulations.

**Table S1.** Total N-glycan coverage of the residues involved in each binding mode using the CHARMM36 description. The results indicate how much of the CD44-HABD surface (that is critical to hyaluronate binding) is covered by N-glycans. For details of the analysis, see Methods above.

| Binding mode | *Realistic asialo:*<br>coverage (%) | *Realistic monosialo:*<br>coverage (%) |
| --- | --- | --- |
| Cryst. | 38 ± 8 | 33 ± 7 |
| Parallel | 36 ± 7 | 36 ± 3 |
| Upright | 24 ± 4 | 20 ± 1 |

### SF. Binding profiles of hyaluronate hexamers in simulations

Figure S6 shows that three initially unbound HA_6_ molecules bind to non-glycosylated CD44 HABD during 20 × 1000 ns trajectories. The occupied epitopes in HABD contacting the HA molecules are at residues 20–26, 38–42, 75–79, 87-100, 105–115, 141–161. These residues match the known epitopes for all three binding modes (14). Qualitatively, we observe only one recognizable binding mode across 20 simulation replicas, with one HA fragment binding to the crystallographic binding mode. Yet, less distinct binding of the HA fragments occurred frequently at distinct binding sites. These observations along with the binding profile agree with the results of the NMR that show simultaneous binding of short HA fragments.

Figure S5 illustrates the probability of different HABD residues to be in contact with the HA hexamers durting the simulations. In panel **a**, the contacts are centered around the R41-centered binding epitope as well as residues in the extended region of HABD. This extended region is also known to undergo major conformational shift from ordered to disordered state upon the binding of ligand. In this region, e.g., arginine R154 have been noted to stabilize the ligand-bound conformation by interacting with the bound ligand in the disordered state of the receptor. In panel **b**), the high probability of contacts in the crystallographic binding groove is due to the other HA hexamer being initially placed into this position. The fact that there was zero dissociation event of the initially-bound HA hexamer clearly shows that the employed simulation force field is capable of reproducing the correct binding interaction.

Figure S6 also shows similar binding behavior for initially unbound HA_6_ molecule that binds to HABD already occupied with another HA_6_ in the crystallographic binding groove. The major binding epitopes locate roughly at residues 38–42, 105–115, 144–169. The C-terminal amino acids, 144–169, correspond to both the upright mode-specific HA binding residues and the residues that are influenced by MEM-85. This observation therefore agrees with the experimental observations and fortifies the notion of a second, lesser affinity binding site at the C-terminal portion of HABD.

**Table S2.** The number of association or dissociation events between individual HA hexamers and HABD during 1000 ns simulations. Data are calculated from simulation set G5 in Table 1 (i.e., HABD with three initially unbound HA hexamers, see Note SB). Association is counted when there are above 50 individual contacts between the atoms of CD44 and HA. Dissociation is counted when number of contacts reaches zero. Weak interactions (i.e., number of contact values between 0 and 50) are omitted.

| Replica | 1 | 2 | 3 | 4 | 5 | 6 | 7 | 8 | 9 | 10 | 11 | 12 | 13 | 14 | 15 | 16 | 17 | 18 | 19 | 20 |
| --- | --- | --- | --- | --- | --- | --- | --- | --- | --- | --- | --- | --- | --- | --- | --- | --- | --- | --- | --- | --- |
| HA<sub>6</sub><sup>1</sup> | 11 | 1 | 1 | 1 | 1 | 9 | 1 | 1 | 6 | 1 | 3 | 7 | 5 | 7 | 3 | 1 | 3 | 5 | 3 | 7 |
| HA<sub>6</sub><sup>2</sup> | 1 | 1 | 3 | 1 | 1 | 5 | 7 | 3 | 4 | 1 | 1 | 2 | 1 | 1 | 1 | 1 | 1 | 1 | 1 | 5 |
| HA<sub>6</sub><sup>3</sup> | 1 | 3 | 11 | 3 | 7 | 2 | 8 | 1 | 5 | 3 | 1 | 1 | 1 | 1 | 3 | 1 | 5 | 9 | 1 | 3 |

Tables S2 and S3 show the number of attachments or detachments between each HA hexamers and CD44-HABD in simulation sets G5 and G6, respectively. One can see that each hexamer attaches to HABD at least once. However, many of the hexamers undergo multiple attachment-detachment cycles during the 1000 ns trajectories, with average number of attachments or detach-ments for the initially-free HA fragments is 3.1 ± 0.5. This indicates that the sampling of the binding surface is adequate. It also suggests that hexamers are most likely too short to readily form any of the possible binding modes at the studied time scales. Namely, hexamers are regarded as the minimum length of HA able to bind CD44 (24). Yet, the only hexamer that was placed into a predefined binding configuration in simulation set G6 remained in this configuration across all simulation repeats. This illustrates that the simulation force field is capable of reproducing the correct binding, thus giving further backing for the other results, too.

**Fig. S3.**
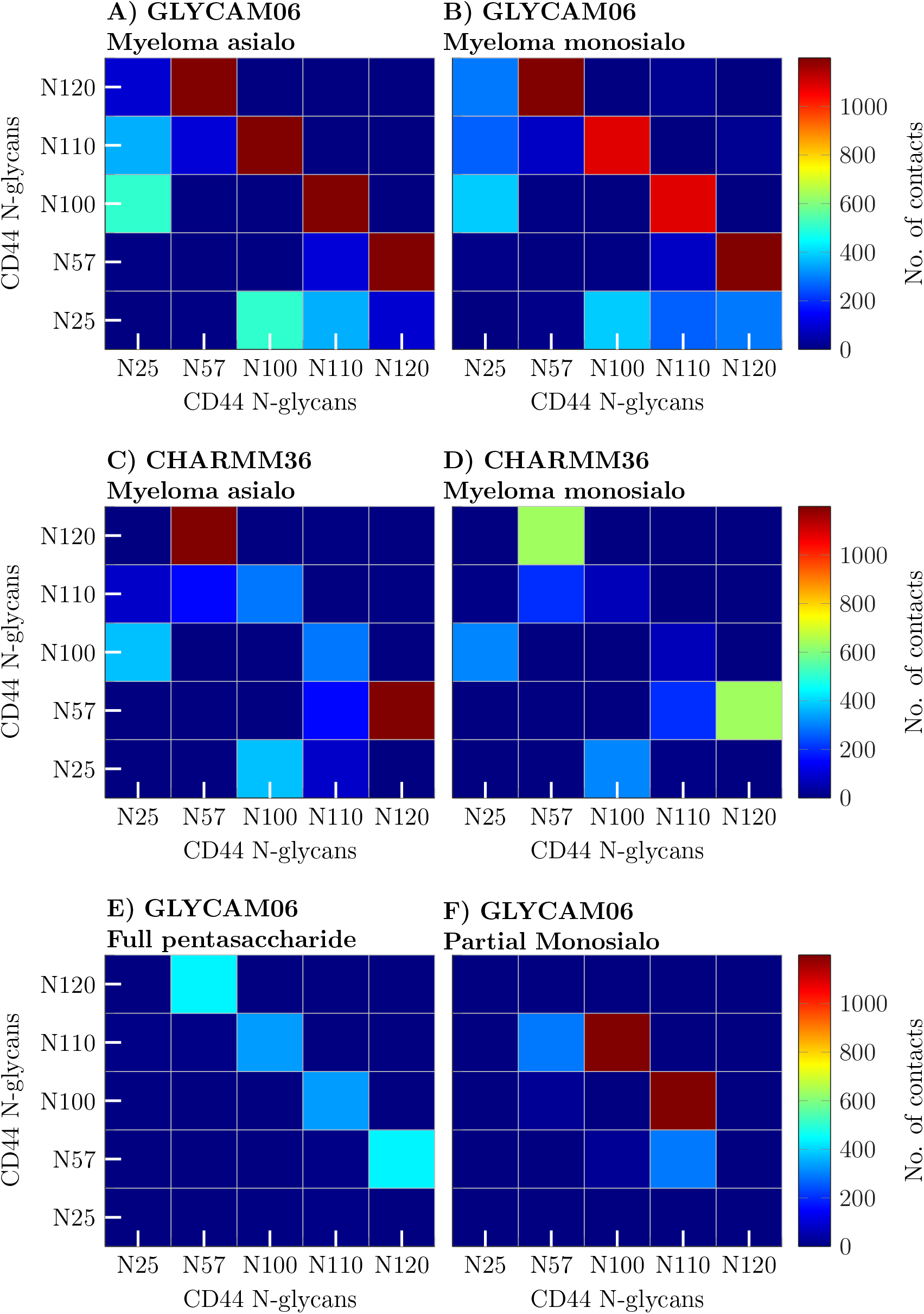
Number of contacts between the five N-glycans on CD44-HABD. Data is time and system averaged from 15 and 3 replicas for the GLYCAM06 and CHARMM36 systems, respectively. In the analysis, all the contacts were summed for each data point and a threshold for a contact was 0.6 nm.

**Fig. S4.**
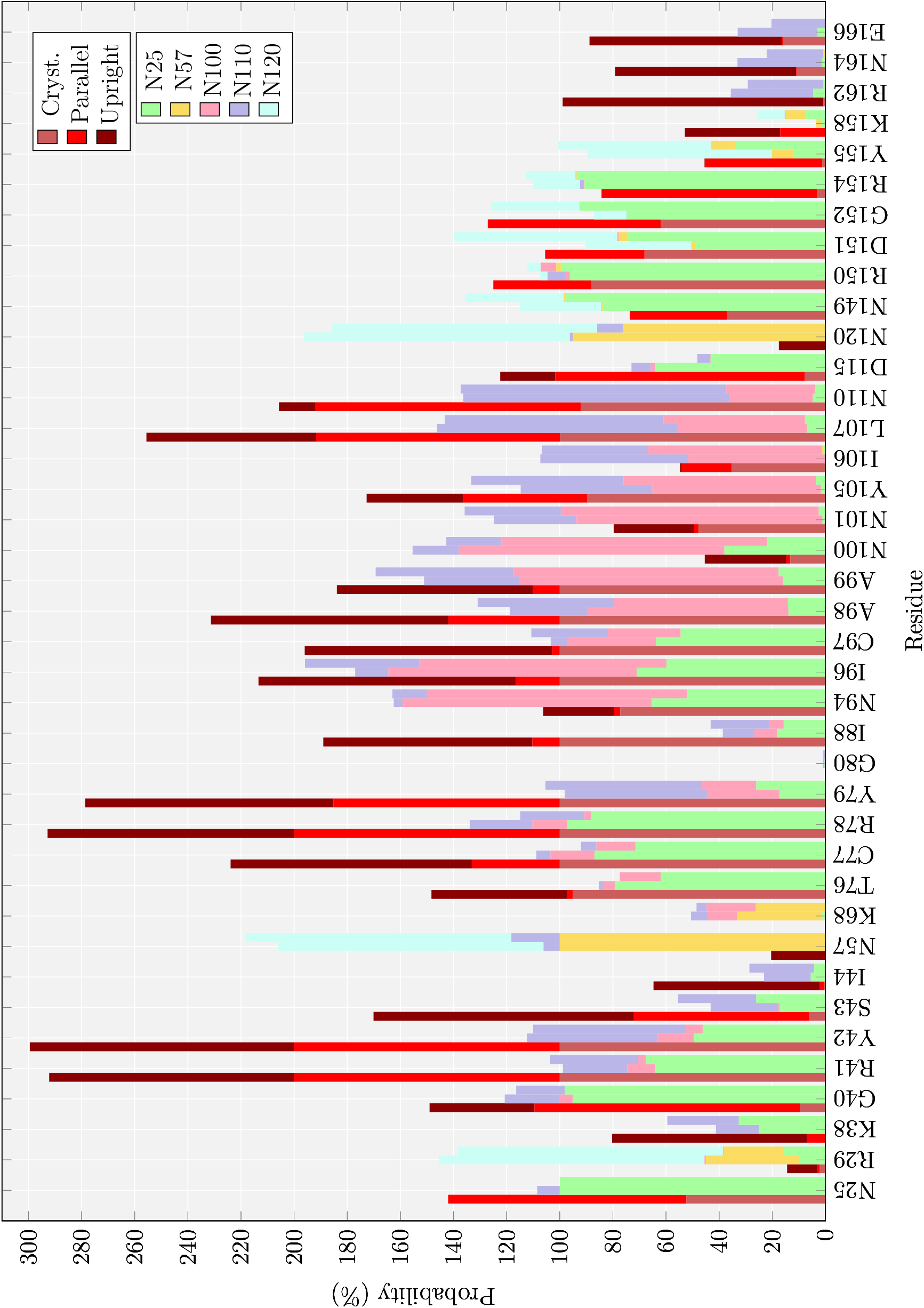
Contact histogram for the observed HA binding residues, according to the literature (14). It shows in how many aggregate simulation frames (%) given binding residue-N-glycan interaction is present. These data are calculated from the *Myeloma monosialo* systems. For comparison, it shows in how many aggregate simulation frames (%) given binding residue-HA interaction is present for each binding mode (Data available in Ref (14)).

**Fig. S5.**
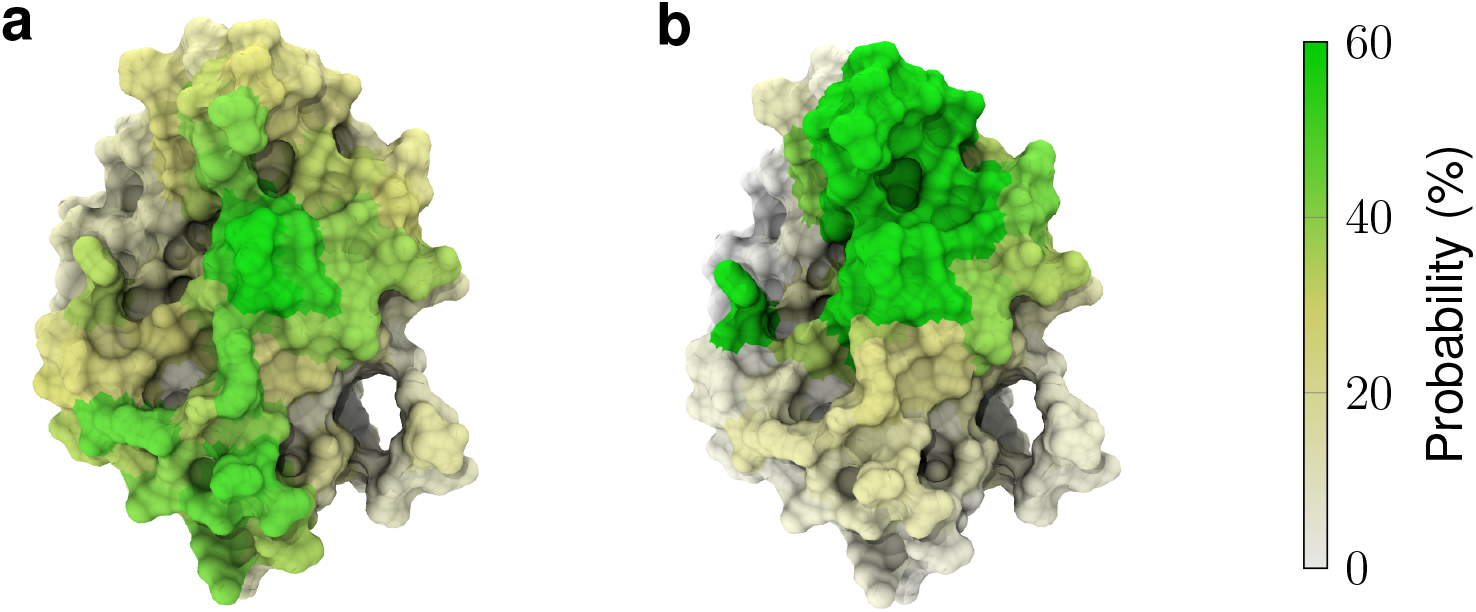
Hyaluronate-perturbed residues in simulations G5 (**a**) and G6 (**b**). The colored surface displays the probability of a given residue to be in contact with HA6 in our simulations.

**Table S3.** The number of association or dissociation events between individual HA hexamers and HABD during 1000 ns simulations. Data are calculated from simulation set G6 in Table 1. Association is counted when there are above 50 individual contacts between the atoms of CD44 and HA. Dissociation is counted when number of contacts reaches zero. Weak interactions (i.e., number of contact values between 0 and 50) are omitted. In these simulations, hexamers number 1 is attached to the crystallographic binding site, and no detachments are recorded. Hexamer number 2 is initially unbound similar to System G5. That is, it can readily sample the surface of HABD for possible binding sites.

| Replica | 1 | 2 | 3 | 4 | 5 | 6 | 7 | 8 | 9 | 10 |
| --- | --- | --- | --- | --- | --- | --- | --- | --- | --- | --- |
| HA<sub>6</sub><sup>1</sup> | 0 | 0 | 0 | 0 | 0 | 0 | 0 | 0 | 0 | 0 |
| HA<sub>6</sub><sup>2</sup> | 5 | 5 | 3 | 4 | 4 | 1 | 0 | 3 | 3 | 2 |

### SG. Example of CD44-hyaluronate binding

Figure S7 shows and example of CD44–hyaluronate binding in both the crystallographic and upright binding modes. As the ligand is depicted at various timestamps, it is easy to see which carbohydrate units interact the most with the protein.

### SH. Spontaneous binding of HA to glycosylated CD44-HABD

Figure S8 shows example structures of observed binding complexes between HA and glycosylated CD44 as well as comparison of HA–CD44 interface area vs HA–N-glycans interface area. These data are calculated from simulations B3–8 (Table 1).

Figure S9 shows relative interaction probability of key arginine residues with HA in glycosylated receptors as compared to non-gkycosylated CD44. The data show the flanking arginines R150, R154, and R162 to have relatively more interactions with the HA ligand in the glycosylated cases. The arginines R41 and R78 at the primary binding site, on the other hand, are less active in binding when the receptor is glycosylated. This illustrates how the sugar shield obstructs the primary binding site.

### SI. Additional observations from the spontaneous binding simulations

Highlighting the sialic acid-induced repulsive effect, HA failed to bind HABD entirely in one replica of the *full polysialo* systems. This is the only occasion with GLYCAM06 force field when we did not observe interactions between CD44 glycoprotein and HA during a 1000 ns trajectory.

The charge-neutral *full extended asialo* glycoform with two galactoses per antennae shows HA binding comparable to the equally-sized but charged *full monosialo* glycoform, which is again significantly lower than that of the shorter monogalactose-terminating *full asialo* glyform. This highlights the role of the size of the N-glycans in masking the binding site and thus determining the HA binding properties.

**Fig. S6.**
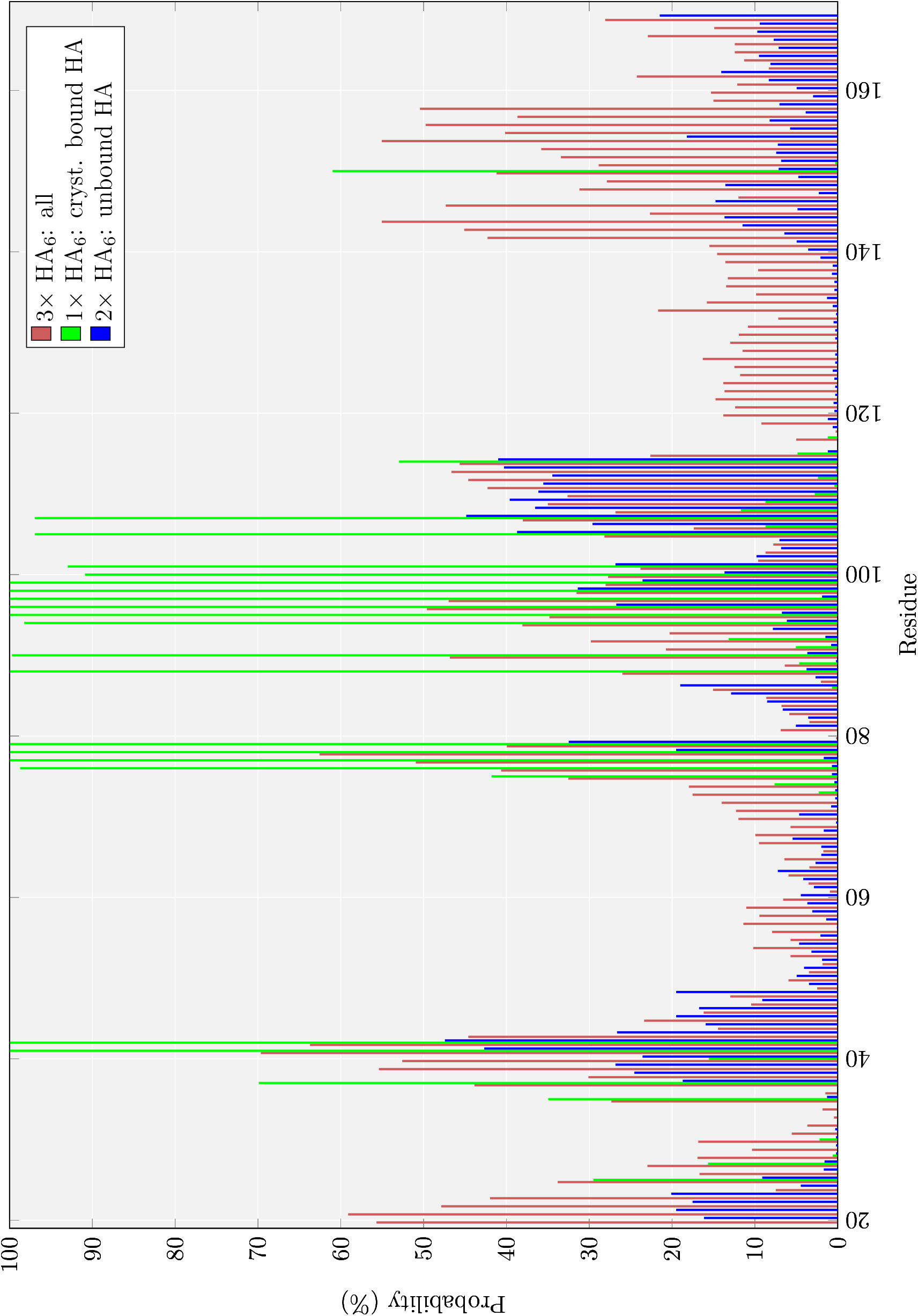
Contact histogram for all HA binding residues vs multiple HA_6_. It shows in how many aggregate simulation frames (%) given binding residue-HA interaction is present. These data are calculated from the G5 and G6 systems in two settings: first, with either three initially unbound HA_6_ molecules, or second, with two HA_6_ molecules from which one is initially bound in the crystallographic mode and one is initially unbound, respectively. The data are calculated and averaged over 20 (system G5) and 10 (system G6) simulation replicas, see Methods for details.

**Fig. S7.**
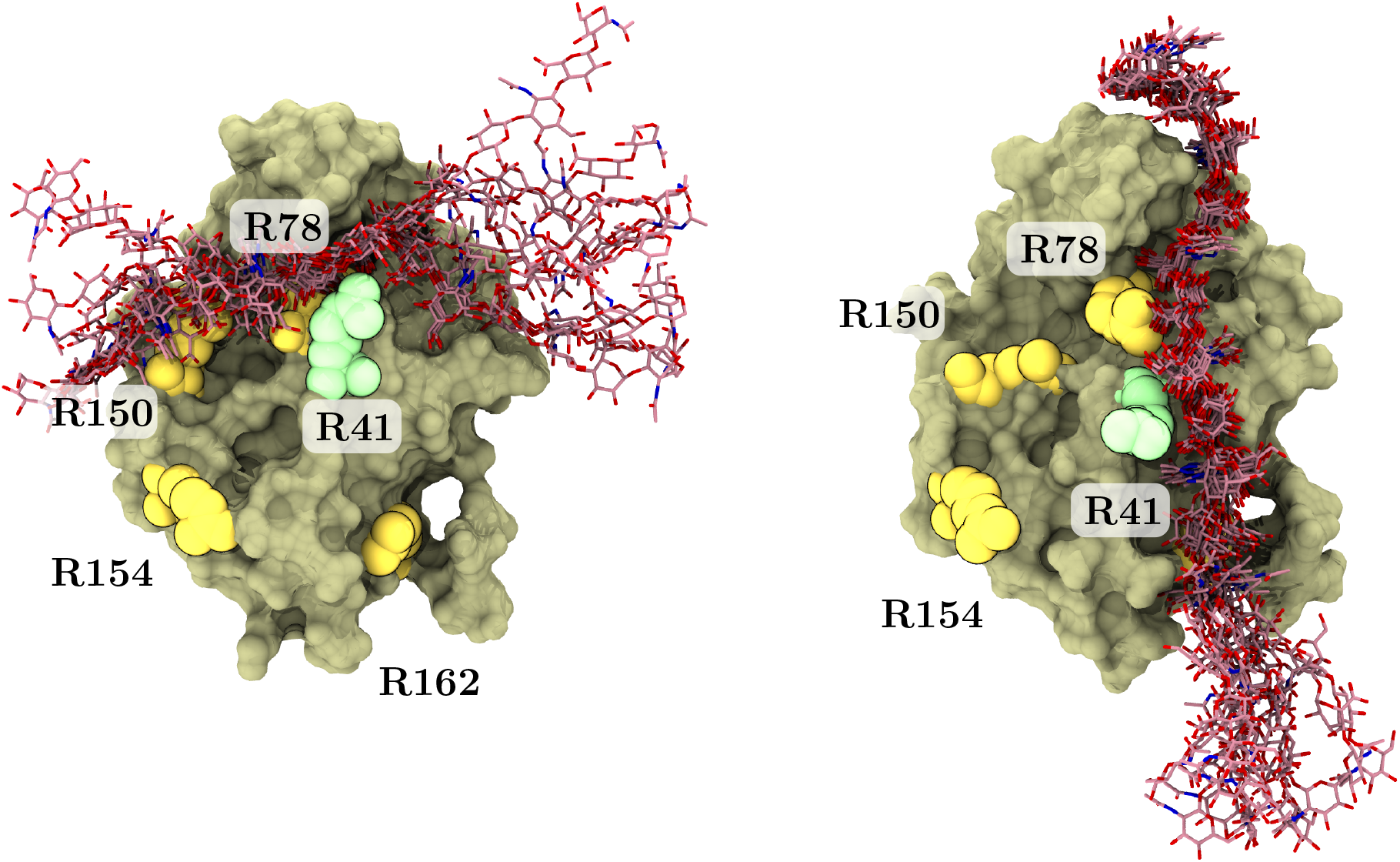
Dynamics of the HA ligand in crystallographic (left) and upright (right) binding modes. The tan surface depicts the protein surface in the first frame of a simulation. The red sticks represent the ligand drawn at every 50 ns into the simulation trajectory. R41 is colored light green. Other key arginines are colored yellow and labeled accordingly. Data are extracted from our previous simulations (14).

**Fig. S8.**
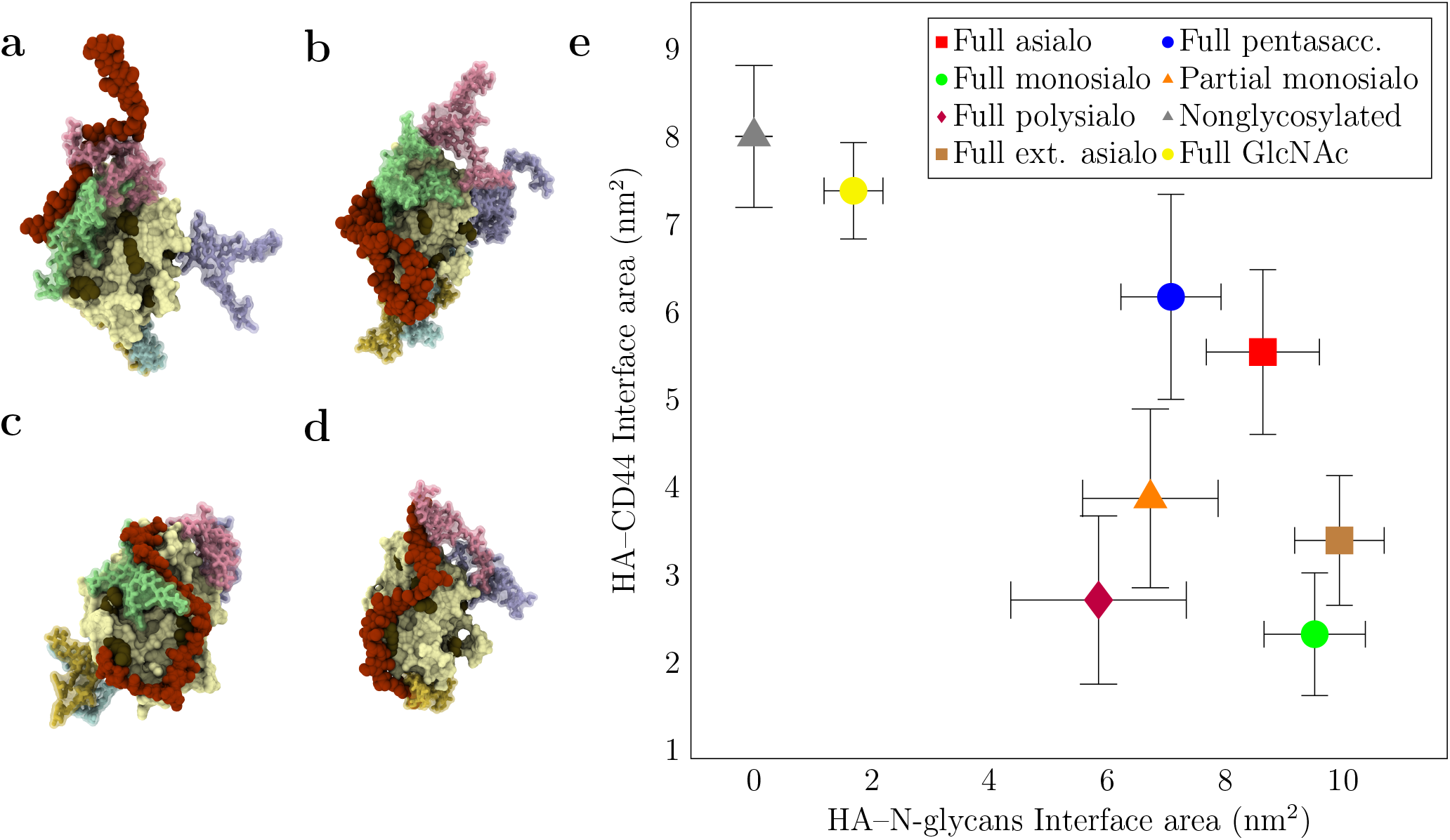
**a-d**: Example structures (*t* = 1000 ns) of HA-CD44 complexes obtained in the spontaneous binding simulations (B3-6 in Table 1). **a**: *full monosialo* glycoform (B4), **c**: *full asialo* glycoform (B3), **b**: *full polysialo* glycoform (B6), and **d**: *partial monosialo* glycoform (B5). **e**: Plot of HA-CD44 interface area vs HA-N-glycans interface area in different glycoforms. The data are calculated with GROMACS tool gmx sasa from systems B1-8. Error bars represent standard errors.

**Fig. S9.**
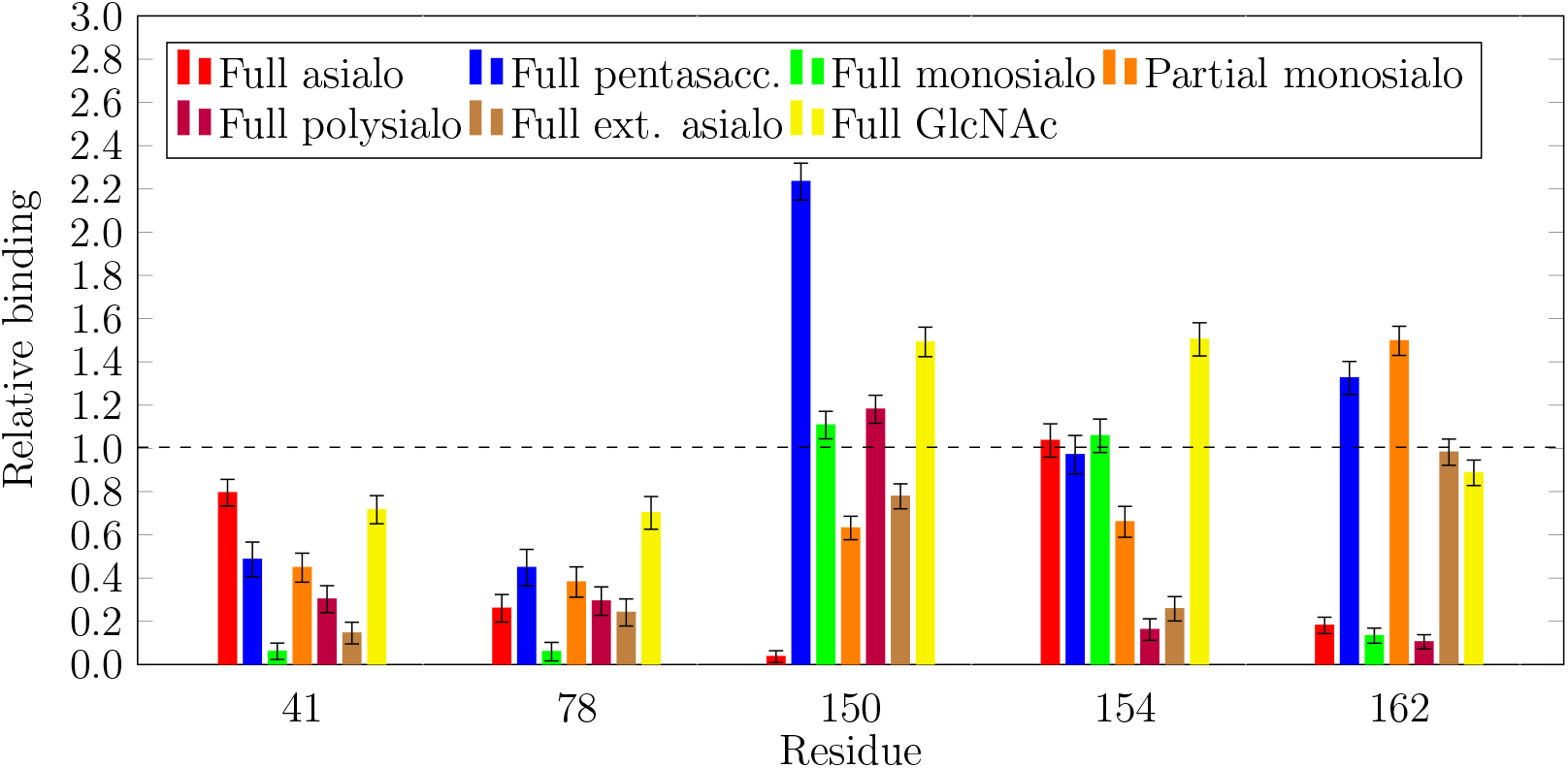
Relative interaction probability of HA with five CD44 arginines in different glycoforms. The data are calculated with GROMACS tool `gmx mindist` by counting the simulation frames in which a contact (HA–Arg minimum distance < *6* Å) is present (excluding the first 200 ns of binding). Values for each glycoforms are then normalized with the reference values from the nonoglycosylated system in order to obtain the relative interaction probabilities. Values above 1 indicate an increase compared to the binding in the non-gkycosylated case, while values below 1 indicate a decrease compared to the non-gkycosylated case.

## Notes

### Competing Interest Statement

The authors have declared no competing interest.

### Summary of Updates

This new version correct issues found in the manuscript during the submission to elife. We replied to the referees' concerns that have been appended to the first biorxiv submission. It is worth mentioning the inclusion of new simulations addressing the hyaluronan binding to glycosylated-CD44 directly.

## Bibliography

1. Anthony P. Corfield and Monica Berry. Glycan variation and evolution in the eukaryotes. Trends in biochemical sciences, 40(7):351–359, 2015. doi: 10.1016/j.tibs.2015.04.004.

2. Rolf Apweiler, Henning Hermjakob, and Nathan Sharon. On the frequency of protein glycosylation, as deduced from analysis of the SWISS-PROT database1. Biochimica et BiophysicaActa (BBA)-General Subjects, 1473(1):4–8, 1999. doi: 10.1016/s0304-4165(99)00165-8.

3. Swagata Halder, Avadhesha Surolia, and Chaitali Mukhopadhyay. Dynamics simulation of soybean agglutinin (SBA) dimer reveals the impact of glycosylation on its enhanced structural stability. Carbohydrate research, 428:8–17, 2016. doi: 10.1016/j.carres.2016.04.009.

4. Xiaolin Huang, Joseph J. Barchi, Feng-Di T. Lung, Peter P. Roller, Peter L. Nara, Jeff Muschik, and Robert R. Garrity. Glycosylation affects both the three-dimensional structure and antibody binding properties of the HIV-1IIIB GP120 peptide RP135. Biochemistry, 36(36):10846–10856, 1997. doi: 10.1021/bi9703655.

5. Najla Arshad, Suhas Ballal, and Sandhya S. Visweswariah. Site-specific N-linked glycosylation of receptor guanylyl cyclase C regulates ligand binding, ligand-mediated activation and interaction with vesicular integral membrane protein 36, VIP36. Journal of Biological Chemistry, 288(6):3907–3917, 2013. doi: 10.1074/jbc.M112.413906.

6. Jonathan W. Lowery, Jose M. Amich, Alex Andonian, and Vicki Rosen. N-linked glycosylation of the bone morphogenetic protein receptor type 2 (BMPR2) enhances ligand binding. Cellular and molecular life sciences, 71(16):3165–3172, 2014. doi: https://doi.org/10.1007/s00018-013-1541-8.

7. Kelley W. Moremen, Michael Tiemeyer, and Alison V. Nairn. Vertebrate protein glycosylation: diversity, synthesis and function. Nature reviews Molecular cell biology, 13(7):448, 2012. doi: 10.1038/nrm3383.

8. Karol Kaszuba, Michał Grzybek, Adam Orłowski, Reinis Danne, Tomasz Róg, Kai Simons, Ünal Coskun, and Ilpo Vattulainen. N-Glycosylation as determinant of epidermal growth factor receptor conformation in membranes. Proceedings of the National Academy of Sciences, page 201503262, 2015. doi: 10.1073/pnas.1503262112.

9. Jayne Lesley, Nicole English, Astrid Perschl, Jennifer Gregoroff, and Robert Hyman. Variant cell lines selected for alterations in the function of the hyaluronan receptor CD44 show differences in glycosylation. The Journal of Experimental Medicine, 182(2):431–437, 1995. doi: 10.1084/jem.182.2.431.

10. Hui Sun Lee, Yifei Qi, and Wonpil Im. Effects of N-glycosylation on protein conformation and dynamics: Protein Data Bank analysis and molecular dynamics simulation study. Scientific reports, 5:8926, 2015. doi: 10.1038/srep08926.

11. Aneta Liwosz, Tianlei Lei, and Maria A. Kukuruzinska. N-glycosylation affects the molecular organization and stability of E-cadherin junctions. Journal of Biological Chemistry, 281(32): 23138–23149, 2006. doi: 10.1074/jbc.m512621200.

12. Yvette Van Kooyk and Gabriel A. Rabinovich. Protein-glycan interactions in the control of innate and adaptive immune responses. Nature immunology, 9(6):593, 2008. doi: 10.1038/ni.f.203.

13. Anne S. van Oosten and Paul A. Janmey. Extremely charged and incredibly soft: physical characterization of the pericellular matrix. Biophysical Journal, 104(5):961, 2013. doi: 10.1016/j.bpj.2013.01.035.

14. Joni Vuorio, Ilpo Vattulainen, and Hector Martinez-Seara. Atomistic fingerprint of hyaluronan-CD44 binding. PLoS computational biology, 13(7):e1005663, 2017. doi: 10.1371/journal.pcbi.1005663.

15. Wolfgang Rudy, Martin Hofmann, Reinhard Schwartz-Albiez, Margot Zöller, Karl-Heinz Heider, Helmut Ponta, and Peter Herrlich. The two major CD44 proteins expressed on a metastatic rat tumor cell line are derived from different splice variants: each one individually suffices to confer metastatic behavior. Cancer Research, 53(6):1262–1268, 1993.

16. Timothy P. Skelton, Chunxun Zeng, Aaron Nocks, and Ivan Stamenkovic. Glycosylation provides both stimulatory and inhibitory effects on cell surface and soluble CD44 binding to hyaluronan. The Journal of Cell Biology, 140(2):431–446, 1998. doi: 10.1083/jcb.140.2.431.

17. Shigeki Katoh, Zhong Zheng, Kenji Oritani, Takaichi Shimozato, and Paul W. Kincade. Glycosylation of CD44 negatively regulates its recognition of hyaluronan. The Journal of Experimental Medicine, 182(2):419–429, 1995. doi: 10.1084/jem.182.2.419.

18. Zhong Zheng, Richard D. Cummings, Philip E. Pummill, and Paul W. Kincade. Growth as a solid tumor or reduced glucose concentrations in culture reversibly induce CD44-mediated hyaluronan recognition by Chinese hamster ovary cells. Journal of Clinical Investigation, 100(5):1217, 1997. doi: 10.1172/jci119635.

19. Nicole M. English, Jayne F. Lesley, and Robert Hyman. Site-specific de-N-glycosylation of CD44 can activate hyaluronan binding, and CD44 activation states show distinct threshold densities for hyaluronan binding. Cancer Research, 58(16):3736–3742, 1998.

20. Alejandro Aruffo, Ivan Stamenkovic, Michael Melnick, Charles B. Underhill, and Brian Seed. CD44 is the principal cell surface receptor for hyaluronate. Cell, 61(7):1303–1313, 1990. doi: 10.1016/0092-8674(90)90694-a.

21. Bryan P. Toole. Hyaluronan: from extracellular glue to pericellular cue. Nature Reviews Cancer, 4(7):528–539, 2004. doi: 10.1038/nrc1391.

22. Helmut Ponta, Larry Sherman, and Peter A. Herrlich. CD44: from adhesion molecules to signalling regulators. Nature Reviews Molecular Cell Biology, 4(1):33–45, 2003. doi: 10.1038/nrm1004.

23. Kayla J. Wolf and Sanjay Kumar. Hyaluronic acid: incorporating the bio into the material. ACS Biomaterials Science & Engineering, 5(8):3753–3765, 2019. doi: 10.1021/acsbiomaterials.8b01268.

24. Peter Teriete, Suneale Banerji, Martin Noble, Charles D. Blundell, Alan J. Wright, Andrew R. Pickford, Edward Lowe, David J. Mahoney, Markku I. Tammi, and Jan D. Kahmann. Structure of the regulatory hyaluronan binding domain in the inflammatory leukocyte homing receptor CD44. Molecular Cell, 13(4):483–496, 2004. doi: 10.1016/s1097-2765(04)00080-2.

25. Suneale Banerji, Alan J. Wright, Martin Noble, David J. Mahoney, Iain D. Campbell, Anthony J. Day, and David G. Jackson. Structures of the Cd44-hyaluronan complex provide insight into a fundamental carbohydrate-protein interaction. Nature Structural & Molecular Biology, 14(3):234–239, 2007. doi: 10.1038/nsmb1201.

26. Huanhuan Han, Martha Stapels, Wantao Ying, Yingqing Yu, Li Tang, Wei Jia, Weibin Chen, Yangjun Zhang, and Xiaohong Qian. Comprehensive characterization of the N-glycosylation status of CD44s by use of multiple mass spectrometry-based techniques. Analytical and bioanalytical chemistry, 404(2):373–388, 2012. doi: 10.1007/s00216-012-6167-4.

27. S. Katoh, S. Maeda, H. Fukuoka, T. Wada, S. Moriya, A. Mori, K. Yamaguchi, S. Senda, and T. Miyagi. A crucial role of sialidase Neu1 in hyaluronan receptor function of CD44 in T helper type 2-mediated airway inflammation of murine acute asthmatic model. Clinical & Experimental Immunology, 161(2):233–241, 2010. doi: 10.1111/j.1365-2249.2010.04165.x.

28. Christina E. Faller and Olgun Guvench. Terminal sialic acids on CD44 N-glycans can block hyaluronan binding by forming competing intramolecular contacts with arginine sidechains. Proteins: Structure, Function, and Bioinformatics, 82(11):3079–3089, 2014. doi: 10.1002/prot.24668.

29. Mitsuhiro Takeda, Hiroaki Terasawa, Masayoshi Sakakura, Yoshiki Yamaguchi, Masahiro Kajiwara, Hiroto Kawashima, Masayuki Miyasaka, and Ichio Shimada. Hyaluronan recognition mode of CD44 revealed by cross-saturation and chemical shift perturbation experiments. Journal of Biological Chemistry, 278(44):43550–43555, 2003. doi: 10.1074/jbc.m308199200.

30. Li-Kai Liu and Barry C. Finzel. Fragment-Based Identification of an Inducible Binding Site on Cell Surface Receptor CD44 for the Design of Protein-Carbohydrate Interaction Inhibitors. Journal of Medicinal Chemistry, 57(6):2714–2725, 2014. doi: 10.1021/jm5000276.

31. Francis W. Jamison II, Theresa J. Foster, Jacob A. Barker, Ronald D. Hills Jr, and Olgun Guvench. Mechanism of binding site conformational switching in the CD44-hyaluronan protein-carbohydrate binding interaction. Journal of Molecular Biology, 406(4):631–647, 2011. doi: 10.1016/j.jmb.2010.12.040.

32. Amanda J. Favreau, Christina E. Faller, and Olgun Guvench. CD44 receptor unfolding enhances binding by freeing basic amino acids to contact carbohydrate ligand. Biophysical Journal, 105(5):1217–1226, 2013. doi: 10.1016/j.bpj.2013.07.041.

33. Jana Škerlová, Vlastimil Král, Michael Kachala, Milan Fábry, Ladislav Bumba, Dmitri I. Svergun, Zdenĕk Tošner, Václav Veverka, and Pavlína Řezáčová. Molecular mechanism for the action of the anti-CD44 monoclonal antibody MEM-85. Journal of structural biology, 191(2): 214–223, 2015. doi: 10.1016/j.jsb.2015.06.005.

34. Jürgen Bajorath, Brad Greenfield, Sandra B. Munro, Anthony J. Day, and Alejandro Aruffo. Identification of CD44 residues important for hyaluronan binding and delineation of the binding site. Journal of Biological Chemistry, 273(1):338–343, 1998. doi: 10.1074/jbc.273.1.338.

35. Brenda M. Sandmaier, Rainer Storb, Kelly L. Bennett, Frederick R. Appelbaum, and Erlinda B. Santos. Epitope specificity of CD44 for monoclonal antibody-dependent facilitation of marrow engraftment in a canine model. Blood, 91(9):3494–3502, 1998. doi: 10.1182/blood.v91.9.3494.3494_3494_3502.

36. Helen M. Berman, John Westbrook, Zukang Feng, Gary Gilliland, T. N. Bhat, Helge Weissig, Ilya N. Shindyalov, and Philip E. Bourne. The Protein Data Bank. Nucleic Acids Research, 28(1):235–242, 2000. doi: 10.1201/9780203911327.ch14.

37. Reinis Danne, Chetan Poojari, Hector Martinez-Seara, Sami Rissanen, Fabio Lolicato, Tomasz Róg, and Ilpo Vattulainen. doGlycans-tools for preparing carbohydrate structures for atomistic simulations of glycoproteins, glycolipids, and carbohydrate polymers for GROMACS. Journal of chemical information and modeling, 57(10):2401–2406, 2017. doi: 10.1021/acs.jcim.7b00237.

38. William Humphrey, Andrew Dalke, and Klaus Schulten. VMD: visual molecular dynamics. Journal of Molecular Graphics, 14(1):33–38, 1996. doi: 10.1016/0263-7855(96)00018-5.

39. Liem X. Dang. Development of nonadditive intermolecular potentials using molecular dynamics: solvation of Li+ and F-ions in polarizable water. The Journal of chemical physics, 96(9):6970–6977, 1992. doi: 10.1063/1.462555.

40. William L. Jorgensen, Jayaraman Chandrasekhar, Jeffry D. Madura, Roger W. Impey, and Michael L. Klein. Comparison of simple potential functions for simulating liquid water. The Journal of chemical physics, 79(2):926–935, 1983. doi: 10.1063/1.445869.

41. Mark James Abraham, Teemu Murtola, Roland Schulz, Szilárd Páll, Jeremy C. Smith, Berk Hess, and Erik Lindahl. GROMACS: High performance molecular simulations through multilevel parallelism from laptops to supercomputers. SoftwareX, 1:19–25, 2015. doi: 10.1016/j.softx.2015.06.001.

42. Berk Hess, Henk Bekker, Herman J. C. Berendsen, and Johannes G. E. M. Fraaije. LINCS: a linear constraint solver for molecular simulations. Journal of Computational Chemistry, 18 (12):1463–1472, 1997. doi: 10.1002/(sici)1096-987x(199709)18:12<1463::aid-jcc4>3.0.co; 2-h.

43. Tom Darden, Darrin York, and Lee Pedersen. Particle mesh Ewald: An N log (N) method for Ewald sums in large systems. The Journal of Chemical Physics, 98(12):10089–10092, 1993. doi: 10.1063/1.464397.

44. Giovanni Bussi, Davide Donadio, and Michele Parrinello. Canonical sampling through velocity rescaling. The Journal of Chemical Physics, 126(1):4101, 2007. doi: 10.1063/1.2408420.

45. Michele Parrinello and Aneesur Rahman. Polymorphic transitions in single crystals: A new molecular dynamics method. Journal of Applied physics, 52(12):7182–7190, 1981. doi: 10.1063/1.328693.

46. Sunhwan Jo, Taehoon Kim, Vidyashankara G. Iyer, and Wonpil Im. CHARMM-GUI: a webbased graphical user interface for CHARMM. Journal of computational chemistry, 29(11): 1859–1865, 2008. doi: 10.1002/jcc.20945.

47. Berk Hess, Carsten Kutzner, David van der Spoel, and Erik Lindahl. GROMACS 4: Algorithms for highly efficient, load-balanced, and scalable molecular simulation. Journal of Chemical Theory and Computation, 4(3):435–447, 2008. doi: 10.1021/ct700301q.

48. Sander Pronk, Szilárd Páll, Roland Schulz, Per Larsson, Pär Bjelkmar, Rossen Apostolov, Michael R. Shirts, Jeremy C. Smith, Peter M. Kasson, David van der Spoel, et al. GRO-MACS 4.5: a high-throughput and highly parallel open source molecular simulation toolkit. Bioinformatics, page btt055, 2013. doi: 10.1093/bioinformatics/btt055.

49. V. Veverka, T. Crabbe, I. Bird, G. Lennie, F. W. Muskett, R. J. Taylor, and M. D. Carr. Structural characterization of the interaction of mTOR with phosphatidic acid and a novel class of inhibitor: compelling evidence for a central role of the FRB domain in small molecule-mediated regulation of mTOR. Oncogene, 27(5):585, 2008. doi: https://doi.org/10.1038/sj.onc.1210693.

50. Ajit Varki, Richard D. Cummings, Markus Aebi, Nicole H. Packer, Peter H. Seeberger, Jeffrey D. Esko, Pamela Stanley, Gerald Hart, Alan Darvill, Taroh Kinoshita, et al. Symbol nomenclature for graphical representations of glycans. Glycobiology, 25(12):1323–1324, 2015. doi: 10.1093/glycob/cwv091.

51. Alicia Lammerts van Bueren and Alisdair B. Boraston. Binding sub-site dissection of a carbohydrate-binding module reveals the contribution of entropy to oligosaccharide recognition at “non-primary” binding subsites. Journal of molecular biology, 340(4):869–879, 2004. doi: 10.1016/j.jmb.2004.05.038.

52. Shigeki Katoh, Taeko Miyagi, Haruko Taniguchi, Yu-ichi Matsubara, Jun-ichi Kadota, Akira Tominaga, Paul W. Kincade, Shigeru Matsukura, and Shigeru Kohno. Cutting edge: an inducible sialidase regulates the hyaluronic acid binding ability of CD44-bearing human monocytes. The Journal of Immunology, 162(9):5058–5061, 1999.

53. Heather C. DeGrendele, Pila Estess, Louis J. Picker, and Mark H. Siegelman. CD44 and its ligand hyaluronate mediate rolling under physiologic flow: a novel lymphocyte-endothelial cell primary adhesion pathway. The Journal of Experimental Medicine, 183(3):1119–1130, 1996. doi: 10.1084/jem.183.3.1119.

54. Heather C. DeGrendele, Pila Estess, and Mark H. Siegelman. Requirement for CD44 in activated T cell extravasation into an inflammatory site. Science, 278(5338):672–675, 1997. doi: 10.1126/science.278.5338.672.

55. Swagata Halder, Avadhesha Surolia, and Chaitali Mukhopadhyay. Dynamics simulation of soybean agglutinin (SBA) dimer reveals the impact of glycosylation on its enhanced structural stability. Carbohydrate research, 428:8–17, 2016. doi: 10.1016/j.carres.2016.04.009.

56. Anirban Polley, Adam Orlowski, Reinis Danne, Andrey A. Gurtovenko, Jorge Bernardino de la Serna, Christian Eggeling, Simon J. Davis, Tomasz Róg, and Ilpo Vattulainen. Glycosylation and lipids working in concert direct CD2 ectodomain orientation and presentation. The Journal of Physical Chemistry Letters, 8(5):1060–1066, 2017. doi: 10.1021/acs.jpclett.6b02824.

57. Diluka Peiris, Alexander F. Spector, Hannah Lomax-Browne, Tayebeh Azimi, Bala Ramesh, Marilena Loizidou, Hazel Welch, and Miriam V. Dwek. Cellular glycosylation affects Herceptin binding and sensitivity of breast cancer cells to doxorubicin and growth factors. Scientific Reports, 7:43006, 2017. doi: 10.1038/srep43006.

58. Pauline M. Rudd, Mark R. Wormald, and Raymond A. Dwek. Sugar-mediated ligand-receptor interactions in the immune system. Trends in biotechnology, 22(10):524–530, 2004. doi: 10.1016/j.tibtech.2004.07.012.

