## Supplementary material for "N-glycosylation blocks and simultaneously fosters different receptor-ligand binding sites: the chameleonic CD44–hyaluronan interaction": Rebuttal elife

This preprint was reviewed using eLife's Preprint Review service, which provides public peer reviews of manuscripts posted on bioRxiv for the benefit of the authors, readers, potential readers, and others interested in our assessment of the work. This review applies only to Version 1 of the preprint: https://www.biorxiv.org/content/10.1101/2020.03.23.004002v1.full

**Summary**

*This manuscript examines how N-linked glycosylation regulates the binding of polysaccharide hyaluronan (HA) to cell surface receptor CD44, to conclude that multiple sites exist but are controlled by the nature of the glycosylation. The reviewers appreciated many aspects of the work, but they have raised serious concerns about the experimental and simulation design. The reviewers suggested that the proposed alternative binding site may not be biologically relevant, as the relevant CD44-HA interactions are multivalent and cannot be supported by that site. They also suggested that the findings are not well supported by the NMR experiments, which could have been extended to allow comparisons of the glycosylation patterns hypothesised. Moreover, the MD simulations, despite being considerable in size, were limited in sampling different possibilities without bias from the initial HA placement, and there is not enough data to convince the readers of thorough sampling and reproducibility.*

We understand the concerns raised in the review process. However, these concerns can be readily explained and fixed, as we explain below and are briefly introduced here.

- Our data are compatible with the currently accepted multivalent interaction of hyaluronan with several CD44 receptors. The argument that our data goes against it stems from an unfortunate figure provided in the first version of the manuscript. This figure suggested that a bound hyaluronan would not be able to span the length the protein in the upright binding mode. That is not true. We now show another, and more relevant snapshot where the bound hyaluronan indeed spans the length of HABD. Hence, we show that multivalent interaction is not precluded by the upright binding mode.
- We also clarify how our extensive simulation data were designed to avoid any bias. We admit that this was not obvious in the phrasing of our previous version.
- Many of the raised issues stem from the lack of certain critical simulations. We have now added these simulations into the revision.

Below we summarize the main issues raised by the reviewers, accompanied by our responses on how we have fixed them in the revised version of the manuscript.

**Reviewer #1**

*1.1. "The authors use MD simulations and NMR to study the cell surface adhesion receptor CD44 with the purpose of understanding the binding of carbohydrate polymer, hyaluronan (HA). In particular, this study focuses on the effects of N-glycosylation of the CD44 glycoprotein on potential HA binding. The authors previously proposed two lower affinity HA binding modes as alternatives to the primary mode seen in the crystal structure of the HA binding domain of CD44, driven by different arginine interactions, but overlapping with glycosylation sites that will affect HA binding. This study suggests that, because the canonical site appears blocked by glycans attached to the surface, HA would instead likely bind to an alternate parallel site with lower affinity, thus changing receptor affinity. The authors do not study HA binding to the glycosylated form directly, but undertake simulations of bound glycans to draw their conclusion. They do, however, place HA near the non-glycosylated CD44 in simulations, although it is not clear that MD sampling has been designed to provide unbiased observations of HA binding, or how the simulations help explain the NMR experiments."*

To better highlight the message, we left out a significant portion of our total simulation data from the initial version of the manuscript. We have now added e.g. simulations of HA binding to the glycosylated form into our revised manuscript. Furthermore, we are confident that our design of the simulation systems allows unbiased sampling of the binding surface. That is, the hyaluronan hexamers were initially placed several nanometres away from the protein surface. After this, they were allowed to spontaneously sample the space and find their respective binding sites during the course of the simulations. They were not placed into the binding sites manually. However, there was a one system with two HA hexamers from which the other was placed into the canonical binding groove. This was done to test where the freely floating hexamer would bind when the primary binding site is taken. These points are illustrated more clearly in the new version of the manuscript. Finally, all our simulation data is publicly available (see the DOIs provided in the paper).

*1.2 "The data rely on libraries of MD simulation, which are substantial, with several replicas of a microsecond each. But what have these simulations really proved with reliability? Figure 2a shows that, while glycans stay roughly where they started, they are dynamic and cover much of the canonical HA binding site, which may be the case. From this the authors imply that the crystallographic site is significantly obstructed, the lower-affinity upright mode remains most accessible, and that the level of occlusion of the main site depends on the degree of glycosylation and size of the oligosaccharides. However, a full simulation of HA binding to this glycosylated surface was not attempted. It would have been good to see the glycans actually block unbiased simulation of canonical binding to the crystallographic site on long timescales (not being dislodged), but allow alternative binding to the parallel site, without initial placement there."*

Commenting both points 1.1 and 1.2, we cropped a large portion of our simulation data from the initial version of the manuscript in order to better highlight the current message. However, we do have extensive simulation data of hyaluronan binding spontaneously to CD44 with different glycosylation patterns. For example, see Figure A below where HA is bound to glycosylated CD44-HABD. These data have been carefully analysed and incorporated into the revised manuscript.

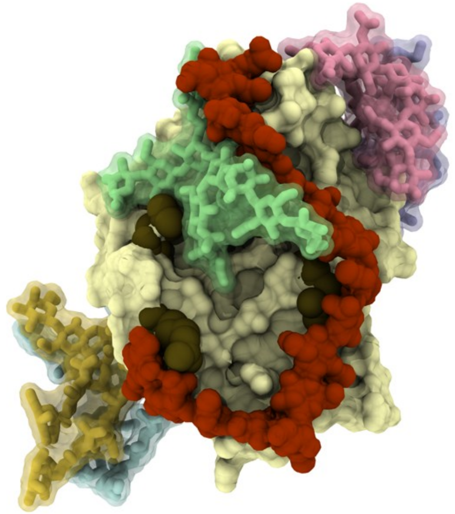

*Figure A. A representative binding pose between HA oligomer (dark red) and glycosylated (light blue, yellow, green, pink and purple) CD44-HABD (pale surface) extracted from our simulations.*

*1.3 "HA was, however, added to the non-glycosylated CD44-HABD surface in simulations, but no clear data is shown to illustrate the extent of sampling, convergence and reproducibility, beyond some statistical analysis of contacts. It seems a total of 30 microseconds of the non-glycosylated protein with 2 or 3 nearby HA placed was run, leading to contacts. But how well did these 30 simulations sample HA movement and relative binding to sites, if at all? Figure 4 suggests that the HA stay where they have been put. As the MD is the dominant source of data for the paper, the extent of sampling and how the outcomes depend on the initial placement of molecules requires proof. Was any sampling of HA movement, such as between canonical and alternative parallel conformations seen in MD?"*

It is important to note that, in the non-glycosylated systems, the hyaluronan hexamers were initially placed several nanometres away from the protein surface. After this, they were allowed to spontaneously sample the space and find their respective binding sites during the course of the simulations. That is, they were not manually placed into the binding sites. We have changed the manuscript to better illustrate this key point.

We have also made the simulation data publicly available (see the DOIs provided in the paper). After inspection of the simulations, we are confident that the reviewers will agree that the results are reliable and do not suffer from convergence problems that could compromise the message we provide.

Moreover, we have even more simulation replicas ready with slightly different initial conditions that provide the same qualitative picture, see Figure B below (compare with Figure 4c in the original submission where one of the hyaluronan hexamers was initially placed in the crystallographic binding site). In these simulations, the hexamers have enhanced contacts with the crystallographic and upright mode residues despite being initially placed far from these binding sites. These simulations were already part of the manuscript.

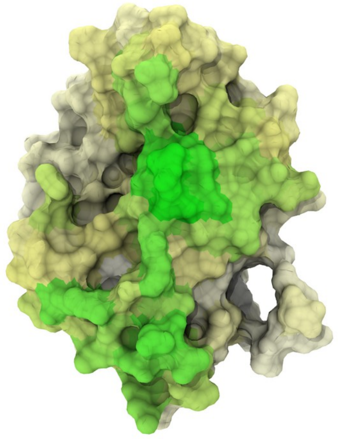

*Figure B. Hyaluronate-perturbed residues in the simulations. The colored surface displays the probability of a given residue to be in contact with HA6 in our additional simulations, where three hyaluronan hexamers were placed in solution far from the binding site.*

*1.4 "The NMR is suggested to show that a short HA hexamer can bind to non-glycosylated CD44-HABD simultaneously in several modes at distinct binding sites, and that MD "correlates" with this. But is this MD biased by initial choices of where and how many HAs are placed, given HA movement is likely not well sampled?"*

The hyaluronan hexamers were initially placed several nanometers away from the binding sites. They were not placed into these binding sites manually. During the simulations the hexamers displayed several binding and unbinding events as they were spontaneously sampling the space and finding their respective binding sites during the course of the simulations.

While we saw multiple binding events to the proposed binding sites, the short size of the hyaluronan fragments was likely not enough for stable binding as the fragments often dissociated within few hundreds of nanoseconds. These points are now more clearly presented in the revised manuscript.

*1.5 “No MD seems to have been used to examine the blocking or lack thereof by antibody MEM-85 in glycosylated or non-glycosylated CD44.”*

This is not feasible using MD simulations, since the structure of the antibody is not available. Fortunately, there is no need for it, as we have direct and reliable experimental evidence using NMR as provided in the manuscript and in our previous work (Skerlova et.al. 2015; doi: 10.1016/j.jsb.2015.06.005). We therefore know where the antibody binds in CD44.

**Reviewer #2**

*2.1 “This manuscript is focused on understanding how N-linked glycosylation regulates the binding of the (very large) polysaccharide hyaluronan (HA) to its major cell surface receptor CD44, a question relevant, for example to the role of CD44 in mediating leukocyte migration in inflammation. The paper concludes that multiple binding sites for HA exist and that their occupancy is determined by the nature of the glycosylation, a suggestion first made by Teriete et al. (2004). The work is based on atomistic simulations with different glycan compositions and NMR spectroscopy on a non-glycosylated CD44 HA-binding domain (HABD) expressed in E. coli. While the question being researched is interesting and of biological relevance, there are flaws in the work.”*

The relevance also stems from the increasing applicability of HA in many biomedical devices and treatment strategies, such as tissue scaffolds and HA-coated nanoparticles for targeted drug delivery. However, we respectfully disagree with the proposed flaws. We address these suggested issues point-by-point in sections 2.2–2.5.

*2.2 “The paper describes how the well-established HA-binding site on CD44 (determined by a co-crystal structure; Banerji et al., 2007) is blocked by N-linked glycosylation (principally at N25 with a contribution from glycans at N100 and N110) and how certain glycans favour binding at a completely distinct binding site that lies perpendicular to the canonical 'crystallographic' binding site. This alternative 'upright' binding site, which has been proposed previously by the authors (Vuorio et al., 2017), needs further supporting experimental data.”*

Indeed, a characterization of the upright mode can be found from (Vuorio et al., 2017. PloS CB. 13:7). This characterization is based on mircoseconds of unbiased MD simulation data as well as extensive free energy calculations. We for example analysed the most important interactions, orientations of the sugar rings, and binding affinities. These data indicate that while the upright binding mode is weaker than the canonical binding mode (Banerji et al., 2007), it has good shape complementarity between the protein, with e.g. most of the sugar rings lying flat on the surface of the protein, indicating that it might have biological relevance.

The supporting experimental data is presented in the current publication. It has been improved and clarified for the revised version of the manuscript.

*2.3 “Firstly, unlike the 'crystallographic' binding site that forms an open-ended shallow groove on the surface of the protein allowing polymeric HA to bind (and multivalent interactions to take place), the 'upright' binding site is closed at one end and can thus only accommodate the reducing end of the polysaccharide (as apparent from Appendix 1 Figure 1). Its configuration means that it would be impossible for this mode of binding to allow multivalent interactions with polymeric HA.*

*This is a major problem since biologically relevant CD44-HA interactions are multivalent where a single HA polymer interacts with a large number of CD44 molecules (e.g. see Wolny et al., 2010 J. Biol. Chem. 285, 30170-30180). So even if this binding site existed, an interaction between a single CD44 molecule on the cell surface with the reducing terminus of an HA polymer would be exceptionally weak."*

We have data to show that our proposed secondary binding mode does not preclude multivalent CD44-hyaluronan interactions. This multivalent interaction, where a long hyaluronan binds simultaneously to several CD44 moieties, is important, and our secondary mode is compatible with it, see the new Figure C below. We acknowledge that our Figure 1 in the Appendix 1 was not sufficiently clear on this matter. That figure illustrated a structure of one possible CD44-hyaluronan complex obtained from just one of our simulations. However, we have a number of related CD44-hyaluronan complexes from other simulations where the bound ligand spans the full length of the protein, showing that the binding site can accommodate more than just the reducing end of the polysaccharide, and this is highlighted in the attached Figure C. Therefore, multivalent binding is not precluded by the upright binding mode. Unfortunately, the figure depicted in the SI of the original manuscript was misleading. To avoid this issue, it has been replaced in the revised manuscript.

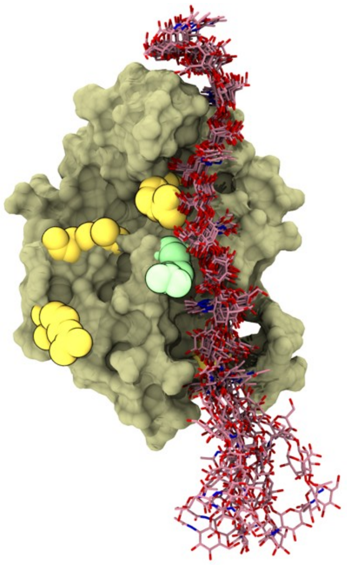

*Figure C. The secondary CD44-hyaluronan binding mode.*

*2.4 "Secondly the NMR experiments performed in this study, purporting to provide evidence for multiple modes of binding, are problematic. Why weren't differentially glycosylated proteins used, i.e. where individual sites were mutated (e.g. +/- N25); this would have allowed comparisons of the glycosylation patterns hypothesised (based on the computer simulations) to favour the 'crystallographic' versus 'upright' modes."*

Indeed, NMR experiments with glycosylated material would be ideal, but obtaining the required quantities of isotopically labelled protein with a **homogeneous** glycosylation pattern is not possible even using the state-of-the-art technology. In addition, the substantially increased molecular weight of the glycosylated protein would be out of the experimental window accessible by NMR spectroscopy. We strongly believe that the message of the paper is already sustained by a combination of our observations based on NMR experiments and MD simulation techniques together with the available literature data as detailed in Appendix A (see below).

While being aware of the difficulties of dealing with glycosylated CD44 using NMR, we designed a way to bypass this issue by combining multiple data from different experimental and simulation setups. All the data support the claims and conclusions made in our paper, see appendix A of this rebuttal. The existence of a weaker binding mode promoted upon glycosylation due to the primary binding site being covered is compatible with all available experimental and simulation data.

*2.4.1 "The CD44 protein used is non-glycosylated, which is not ideal when testing a hypothesis that glycosylation patterns regulate the binding site. While making homogenous glycoproteins is not without its challenges, the authors should consider whether advances in bacterial glycoengineering could be used to generate suitable reagents for experimental analysis; while the glycosylated CD44 construct will be of higher molecular weight (~9-12 kDa) it should not preclude HSQC type shift mapping experiments."*

The bacterial production of the HABD is followed by a complex redox-dependent refolding. To our knowledge, there is no example of refolding of a glycosylated protein reported to date, as the presence of a glycosylation forms an additional barrier for efficient refolding. Even, if this would be successful, the 'dragging' force of oligosaccharide chains would significantly exceed its nominal molecular weight, which would lead to excessive relaxation-driven line broadening.

*2.4.2 "Regarding interpretation of NMR experiments, the chemical shift values result from the sum total of local chemical environment. Perturbations arise from changes in protein environment, which includes both direct contact with a bound ligand and ligand-induced conformational changes. Thus, it is impossible to distinguish from chemical shift perturbation data alone whether perturbations are due to proximity to a bound ligand, ligand-induced conformational changes or a mixture of both. The magnitude and pattern of perturbations therefore cannot be simply interpreted to delineate different binding sites. The observed exchange regimes for a nucleus depend on the magnitude of its chemical shift perturbation, which likewise depends on both its proximity to the ligand and ligand-induced conformational changes in a complex manner. Indeed, one expects to see chemical shift perturbations with different exchange regimes in a single ligand-binding process where there is a large protein conformational change occurring on binding. Therefore, the different exchange regimes cannot be simply interpreted to delineate different binding sites. The comment by the authors "we also observed different exchange regimes that allowed us to discriminate between different HA binding sites" was not found to be convincing."*

We do believe we already take into account the different exchange regimes in the manuscript (see below).

*2.5 "Furthermore, previous NMR studies have shown that the binding of HA to CD44 causes a considerable number of chemical shift changes due to the induction of a large conformational change in the protein (Teriete et al., 2004; Banerji et al., 2007), making it very difficult to identify amino acids directly involved in HA binding based on the NMR data. Moreover, this conformational change has been fully characterised for mouse CD44 with structures available in the absence and presence of HA (Banerji et al., 2007); this information should have been used to inform the interpretation of the shift mapping. In fact, the way in which the shift mapping data are interpreted is simplistic and doesn't fully take account of the reasons that NMR spectra can exhibit different exchange regimes."*

We interpreted the NMR data very carefully. We are aware of the extent of conformational changes induced by HA binding in CD44-HABD, in fact, we identified them as a molecular mechanism underlying the mode of action for the MEM-85 antibody (Skerlova et.al. 2015; doi: 10.1016/j.jsb.2015.06.005). Therefore, we focused on the differential changes in the NMR signal positions of surface exposed residues upon titration with HA and MEM-85. We also observed different exchange regimes that allowed us to discriminate between different HA binding sites. We emphasized these points in the revised manuscript.

*2.5.1 "Regarding site-directed mutagenesis. K38 and R162 were indicated to be key residues specific for the 'secondary binding site', but data available in the literature (on glycosylated CD44) do not support their involvement in HA binding. The authors would need convincing data showing that mutations (that did not perturb the fold) could differentiate the different modes of binding, "*

In fact, there is literature data available on both of these residues playing a role in the HA binding. For example, (Bajorath et al., 1998) uses mutagenesis to find K38 to be important for the binding without disturbing the fold. Furthermore, (Peach et al., 1993) shows that mutating R162 to alanine decreases the binding.

*2.5.2 "Regarding the simultaneous binding of 2 HA oligosaccharides to non-glycosylated CD44. NMR titration data should be interpreted with caution due to the large conformational perturbation that occurs on binding. The interaction of HA with essentially the same non-glycosylated CD44 construct as used in the submitted paper has been carried out previously by ITC (at very high CD44 and HA concentrations). This showed a stoichiometry of 1 (Banerji et al., 2007. Nat Struct Mol Biol. 14:234); being inconsistent with simultaneous binding of multiple HA oligosaccharides to a single CD44 molecule."*

We are aware of the extent of conformational changes induced by HA binding in CD44-HABD, in fact, we identified them as a molecular mechanism underlying the mode of action for the MEM-85 antibody (Skerlova et.al. 2015; doi: 10.1016/j.jsb.2015.06.005). Thus, we made use of MEM-85 and looked at differences with and without the antibody.

**Reviewer #3**

*Vuorio and colleagues combine atomic resolution molecular dynamics simulations and NMR experiments to probe how glycosylation can bias binding of hyaluronan to one of several binding sites/modes on the CD44 hyaluronan binding domain. The results are of interest specifically to the field of CD44 biophysics and more generally to the broad field of glycosylation-dependent protein-ligand binding. The manuscript is clearly written, and the combination of data from computational and experimental methodologies is convincing. I especially commend the authors on the thorough molecular dynamics work, wherein they ran multiple simulations at microsecond timescale and tried different force fields to minimize the likelihood of their findings being an artifact of a particular force field.*

The use of multiple force fields was indeed meant to alleviate potential force field specific issues. Likewise, the use of multiple simulation repeats with different starting positions and randomized atom velocities were meant to provide comprehensive statistics, minimizing the chances of over-interpreting any isolated phenomena.

**Appendix A**: Summary of the logic of the research procedure together with the experimental, simulation and literature results supporting each step.

1. Non-glycosylated CD44 binds HA *(NMR experiments)*
2. Non-glycosylated CD44 also binds HA in the presence of MEM-85 *(NMR experiments)*
3. Glycosylated CD44s that bind HA do not bind HA in the presence of MEM-85 *(from literature [J. Bajorath, B. Greenfield, S. B. Munro, A. J. Day, A. Aruffo, Journal of Biological Chemistry 273, 338 (1998).]).*

4. We show the MEM-85 binding site in non-glycosylated CD44 to be far from the canonical crystallographic binding region (NMR experiments). This MEM-85 binding site region is mostly inaccessible to typical N-glycans found in CD44 (MD simulation). Therefore, we expect that MEM-85 binds glycosylated CD44 in the same region. *(Our working hypothesis)*

5. Taken together, the above points indicate that MEM-85 covers at least partially the relevant HA binding mode in glycosylated CD44, which has zero overlap with the crystallographic mode. This supports the idea of an alternative binding mode to the crystallographic mode which must be readily available for glycosylated CD44. *(Our finding)*

6. Furthermore, heavily glycosylated CD44 variants cover a significant fraction of the crystallographic mode binding region (MD simulation), potentially making it unavailable for HA binding. This explains why non-glycosylated CD44 binds HA in the presence of MEM-85 (i.e., crystallographic mode is free), while glycosylated CD44 does not (i.e., crystallographic mode is covered with N-glycans). The upright region, on the other hand, experiences only minor coverage by the N-glycans in the glycosylated CD44 and is thus free to bind the ligand *(MD simulations)*.

7. Non-glycosylated CD44 binds HA simultaneously with the crystallographic mode and the upright mode when exposed to high concentrations of small hyaluronan hexamers *(NMR titration and MD simulations)*.

8. Pinpointing the position of the residues that experience the largest chemical shift during the titration experiments using non-glycosylated CD44 clearly shows the fingerprint of the canonical crystallographic mode but also a region compatible with our proposed upright mode *(NMR titration experiments)*. These results are compatible with our simulations of several hyaluronan hexamers *(MD simulation)*.

9. Upright binding mode is accessible to hyaluronan binding in the glycosylated CD44 *(MD simulations shown in this letter that could be included to the paper if deemed necessary)*.

Glycosylation, and glycoscience in general, is one of the most challenging topics to understand in life sciences. We believe that our paper makes a very significant contribution to this area of research in the context of a central research problem and is exceptionally able to provide an atomic-level description of the HA-CD44 interaction under unambiguously known conditions.

Yours sincerely,
Dr. Hector Martinez-Seara
